# Region-specific Brain Targets Drive Circuit Formation and Maturation of Human Retinal Ganglion Cells

**DOI:** 10.1101/2025.05.10.651091

**Authors:** Kang-Chieh Huang, Eyad Shihabeddin, Han-Yin Jeng, Qudrat Abdulwahab, Victoria R. Cuevas, Anthony Ho, Claire Young, Melody Hernandez, Justin Dhindsa, Mikhail Y. Kochukov, Snigdha Srivastava, Benjamin Arenkiel, Jason S. Meyer, Nicholas M. Tran, Melanie A. Samuel

## Abstract

Human vision relies on retinal ganglion cells (RGCs), and their connectivity with distinct brain regions enables higher order visual processing. RGCs vary considerably between species, and many model organisms display distinct RGC types and innervation patterns from those in humans. There is thus a need for robust models of human RGC circuit formation that preserves innervation specificity. Here, we developed an *in vitro* microfluidics eye-to-brain connectivity model using human pluripotent stem cell (hPSC)-derived RGCs to assess brain region-specific connectivity features. We found that cultured human RGCs segregate their dendrites and axons and display axonal features that align with those of their *in vivo* human RGC counterparts. To identify brain target surrogates, we conducted bioinformatic similarity assessments, which revealed high conservation and lower variability between human LGN and mouse counterparts relative to available thalamocortical organoids. We thus provided region-specific mouse visual targets to assess specificity and activity. We found that human RGC axons terminals differentially connected with distinct mouse retinorecipient brain regions and promoted region-specific neural activity. Increased synapse formation occurred between RGCs and lateral geniculate neurons relative to that with suprachiasmatic nucleus neurons, modeling retina brain connectivity differences. Both retinorecipient partners induced the formation of more synapses relative to non-retinorecipient brain target controls. These results suggest that human RGC innervation properties are preserved in culture systems and that human RGCs can differentially sense and respond to retinorecipient targets to control wiring outcomes. These systems may aid in the discovery of wiring factors for potential therapeutic applications.

**Significance Statement:** This study presents a novel *in vitro* model to investigate human retinal ganglion cell (RGC) connectivity, using human pluripotent stem cell-derived RGCs and specific mouse brain targets. By modeling eye-to-brain connections in a microfluidics device, we reveal that cultured human RGCs can form selective, region-specific synapses with mouse-derived brain areas like the lateral geniculate nucleus and the suprachiasmatic nucleus. The findings demonstrate that human RGCs retain their innervation specificity presences in culture, mimicking *in vivo* human connectivity patterns. This model thus provides a powerful tool for understanding the factors controlling human-specific brain wiring, with potential applications in therapies for visual and neurological disorders.

## Introduction

Retinal ganglion cells (RGCs) are vital conduits for transmitting visual information from the eye to the brain, and their precise connectivity with retinorecipient brain regions is essential for proper visual processing^1,2^. To establish region-specific connectivity patterns, RGC axons traverse long distances and navigate through various brain circuits. This process involves extending axons within the retina, reaching the optic disc, forming the optic nerve, and projecting either ipsilaterally or contralaterally to synapse with distinct postsynaptic targets.^3^ These connectivity patterns differ across brain regions, with some areas receiving dense RGC innervation while others receive sparse inputs from defined RGC types. However, the mechanisms governing RGC axon connectivity upon arrival to a target region remain largely unclear. Two primary models have been proposed to explain this process: the permissive model, where simply reaching a target region is sufficient to promote connectivity, and the instructive model, where RGCs actively respond to region-specific cues to establish appropriate synapses^2,4,5^. Distinguishing between these alternative models is a critical step toward identifying the molecular signals driving RGC brain innervation.

Understanding how specific postsynaptic regions influence RGC maturation and connectivity is crucial for two main reasons. First, defects in RGC connectivity with their post-synaptic targets contribute to many human-specific RGC-related visual disorders, including glaucoma^6,7^. Second, significant differences in RGC diversity and connectivity between humans and mice highlight the necessity of human-specific RGC models^3^. While mice exhibit approximately 47 RGC subtypes^8,9^, primates have roughly half that number, with certain RGC types, such as midget cells, dominating the population in primates^10,11^. The abundance of each RGC type also varies between species, with one type of RGC called midget ganglion cells accounting for ∼70% of all primate RGCs^12,13^, while its mouse homolog accounts for only 2% of all mouse RGCs^14^. This emphasizes the need to investigate human-specific RGC development and their synaptic connectivity patterns.

Human pluripotent stem cells (hPSCs) have enabled the generation of retinal organoids containing human RGCs (hRGCs), providing a promising model system for human retina visual circuits. However, hRGCs in these systems often lack polarization, fail to extend long axons, and eventually die^15,16^. This death may be due in part to the lack of trophic support from postsynaptic RGC partners^16^. To address these challenges, we previously developed two- and three-dimensional co-culture systems combining retinal organoids with thalamic cultures, demonstrating that hRGC axons can grow and that their survival is enhanced in the presence of innervation targets^16^. Despite these advances, key questions remain: Do hRGCs *in vitro* develop cellular polarization and structural features characteristic of maturing RGCs *in vivo*? And do hRGCs exhibit the same preference for specific retinorecipient targets, such as the lateral geniculate nucleus (LGN) over the suprachiasmatic nucleus (SCN), observed *in vivo*^17,18^? Finally, how does RGC synapse formation with brain neurons influence the reciprocal maturation of both hRGC and their postsynaptic targets?

To address these gaps, we developed a human eye-to-brain connectivity platform that supports hRGC polarization and integrates region-specific postsynaptic targets, enabling the study of synapse formation and circuit development of human retinal ganglion cells. Using hPSCs from two independent lines, we generated retinal organoids and purified hRGCs, which were cultured in a microfluidics system. This approach facilitated the development of polarized dendrites and axons in hRGCs, including the formation of axon initial segments with *in vivo*-like molecular composition and organization. hRGCs displayed diverse dendritic morphologies, consistent with advanced maturation. When hRGC axons were cultured with mouse-derived retinorecipient targets, such as the LGN and SCN, hRGCs exhibited increased synapse formation, as evidenced by synapse quantification and circuit tracing. Notably, hRGCs formed more synapses with the LGN versus SCN targets, modeling *in vivo* connectivity patterns, and both the LGN and SCN were favored over non-retinorecipient controls. Together, these findings demonstrate that our *in vitro* eye-to-brain connectivity model recapitulates *in vivo* region-specific innervation and suggest that conserved, cell-intrinsic mechanisms drive human RGC connectivity with specific brain regions. This model provides a valuable tool for advancing our understanding of human RGC development and has implications for restoring connectivity in visual disorders and guiding the integration of transplanted cells.

## Results

### hPSC-derived RGCs develop polarized dendrites and axons

Shortly after their birth, RGCs extend local dendrites that remain near the soma and develop long axons that exit the eye, project through the optic chasm, and establish connections with specific post-synaptic targets in different brain regions^3^. To determine if hRGCs were capable of this distinctive dendritic and axonal polarization, we generated hRGCs from three-dimensional retinal organoids (**Supplemental Figure 1A**). We used two hPSC lines that genetically encoded an RGC reporter: the human embryonic stem cell (hESC) H7, which encodes a tdTomato-Thy1.2 reporter (**Figure 1A)**, and the human induced pluripotent stem cell (hiPSC) WTC11, which expresses Thy1.2 under the control of the RGC-specific transcription factor Brn3b^19^. In both lines, the spatial and temporal generation of retinal neuron subtypes modeled the developmental stages observed *in vivo*^20–23^, with cycling CHX10 and Ki-67 positive retinal progenitors visible in the outer retina early in development, and laminated, fate committed neurons present later in organoid development (**Supplemental Figure 1B-D**). Human RGCs are among the earliest generated retinal neurons and are born beginning at fetal week four^24^. Similarly, hRGCs (BRN3+) were present in human retinal organoids by culture day 30. The ratio of hRGCs to retinal progenitor cells (CHX10+) increased over time and hRGCs became restricted within the inner retina layer (**Supplemental Figure 1C-D**).

**Figure 1:**
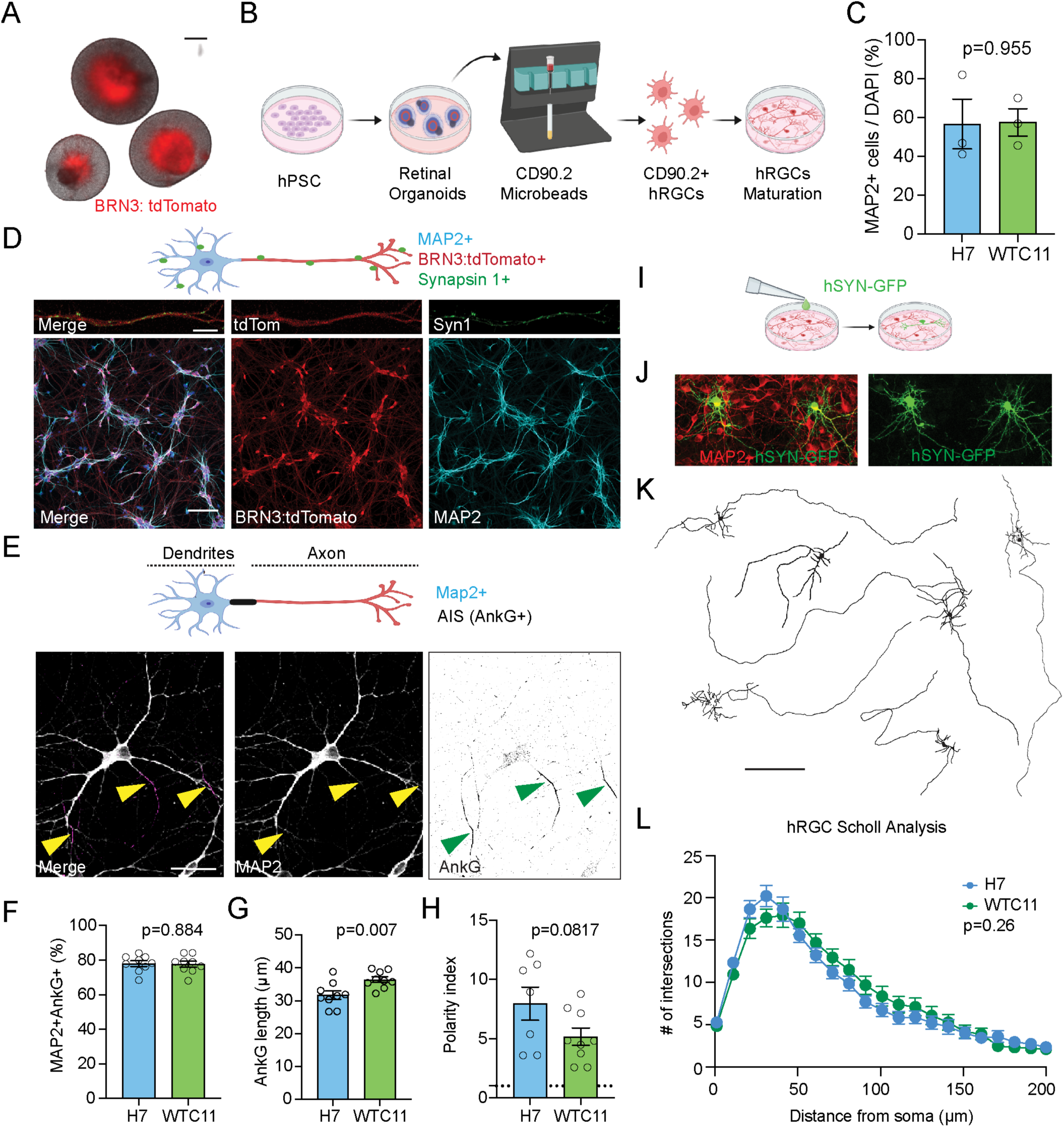
hRGCs mature and polarize in long term cultures. **A.** hRGCs can be identified in the inner retinal layers of organoids generated from the H7 tdTomato-Thy1.2 line. Scale bar = 200 µm.**B**. Schematic of hRGC generation and isolation using CD90 (Thy1.2) conjugated magnetic beads. This approach can be used to purify hRGCs from both hESC line H7, which carries tdTomato-Thy1.2, and the iPSC line WTC11, which carries Thy1.2 alone under the transcription factor BRN3. **C.** Quantification of hRGC purity from both the WTC11 and the H7 iPSC lines. n = 3 biological replicates cultures and >200 cells were count per biological replicates from both lines. **D**. Representative images of hRGCs at three weeks post plating. hRGCs (tdTomato, red) express high levels of the dendrite marker MAP2 (cyan) clustered around the cell somas. Scale bar = 100µm. Representative images of hRGCs following staining for the synapse marker Syn 1 (green) at three weeks post plating. Scale bar = 10µm. **E**. Representative images of hRGCs following staining for the dendrite marker MAP2 (grey) and the axon initial segment marker Ankyrin G (AnkG, magenta). At three weeks post plating, hRGCs show distinct axon initial segments (AISs), which appear close the soma (arrows). Scale bars = 100µm. **F.** Quantification of hRGCs derived from both hPSC lines show that the majority of cells possess an AIS. There is no significant difference the number of neurons with an AIS between the two lines. **G**. Quantification of AIS length in hRGCs following staining with AnkG. In both cell lines, the AIS is 30-40µm long, with those in WTC11 derived hRGCs appearing slightly longer. **H**. The polarity index was calculated as the intensity ratio of MAP2 and AnkG. Both cell lines exhibited strong polarization, with polarity index values ranging from 2 to 4, and no significant differences were observed between the hRGC groups (n = 9 images from 3 biological replications, each image contains >20 hRGCs). **I**. Schematic of sparse labeling hRGCs using lipofectamine transfection. **J.** Representative images of hRGCs a week after hSYN-GFP sparse labeling. Scale bar = 50µm **K.** Representative images of single hRGC neurite reconstruction. Scale bar = 1000µm. **L**. Sholl analysis revealed that most hRGCs displayed a high density of neurites near the soma, consistent with dendrite and axon segregation. No significant differences were detected between hRGCs derived from the two hPSC lines. n = 18-19 hRGCs from 3 biological replicates in both hPSC lines. Data are presented as average values ± the s.e.m.

We next examined hRGC neural outgrowth, synapse formation, polarization, and structural complexity of purified RGC cultures over time to assess their maturation. To purify hRGCs, we performed enzymatic digestion followed by tagging and isolation using an antibody to the RGC-specific surface protein CD90.2 (Thy1.2) coupled to magnetic beads (**Figure 1B**). These generated cultures enriched for hRGCs for studies of their development and organization (**Figure 1C-D**). Staining for the dendritic marker MAP2^25^, the synapsin marker Syn1, and the axon initial segment (AIS) marker ankyrin G (AnkG)^26^ revealed that three weeks post plating, RGCs showed segregated dendrites and axons, with 80% of RGCs showing a defined AIS (**Figure 1E-F**). The percent of hRGCs with AISs did not differ significantly between the hPSC lines (*p = 0.884,* **Figure 1F**). AIS structures were also enriched for β-spectrin, a cytoskeletal protein concentrated at the AIS, and for voltage gated sodium channels, which are anchored and clustered at the AIS by βIV-spectrin (**Supplemental Figure 2A**).

To test whether AIS complexity, organization, and structure were conserved in human RGCs, we measured AIS length and distribution. hRGC AISs were between ∼30 and 40 um long, with those derived from WTC11 iPSCs showing a small but significant increase in AIS length compared to H7 hPSC-derived hRGCs (*p = 0.007,* **Figure 1G**). These values are similar to those reported in mouse RGCs^27^ as well human neuronal AIS structures *in vivo*^28^ and *in vitro*^29^. To assess whether neuronal processes were specializing into distinct axons or dendrites necessary for functional circuitry, we calculated the hRGC polarity index^30,31^. The polarity index is defined as the ratio of AIS immunofluorescence (AnkG) in the axon to AIS immunofluorescence in the dendrite. Values greater than 1 are indicative of polarization as they show that the AIS is enriched in neurites that do not contain dendritic markers^32^. hRGCs from both cell lines were highly polarized, with polarity index measures between 2 and 14, and there was no significant difference in polarity between the hRGC groups (*p = 0.817,* **Figure 1H**). We then examined the shape and structure of individual hRGCs via sparse transduction with a hSyn1-GFP vector and single cell reconstruction (**Figure 1I-K**). The majority of hRGCs had dendrites that clustered around the soma, and a single long axon up to a few millimeters in length (**Figure 1K**). Sholl analysis showed that hRGCs from both lines have a similar level of neurite complexity, with increased dendritic branching near the soma (**Figure 1L**). Together, these data indicate that hRGCs polarize, mature, and develop conserved molecular, structural, and morphological features reflective of human RGCs.

### hRGC polarization is maintained in a microfluidics system

To examine the impact of post-synaptic targets on hRGC connectivity, we first needed to devise a system in which hRGC axons can be spatially separated from their dendrites. Toward this goal, we utilized a microfluidics system in which hRGC cell bodies are plated in the somatic compartment and the axons are allowed to grow into the distal axonal compartment through microgrooves connecting the chambers (**Figure 2A-B**)^16^. First, we tested the ability of hRGCs to develop and maintain polarization in the microfluidics system. Purified H7 or WTC11-derived hRGCs were seeded into the somatic compartment, and additional brain-derived neurotropic factor (BDNF) and ciliary neurotrophic factor (CNTF, see Material and Methods) were added to the axonal chamber to enhance RGC axon extension into the contralateral axonal compartment^16^. We found that hRGC dendrites remained restricted to the somatic chamber, as indicated by localization of 99.33% MAP2 positive processes confined to this region (**Figure 2C-E**). In contrast, histological labelling showed that 62.86% of total fluoresce for the axon marker SMI-312 was in the somatic chamber, 3.61% was in the microgrooves, and 33.53% was in the axonal chamber of the microfluidics devices (**Figure 2C-E)**.

**Figure 2.**
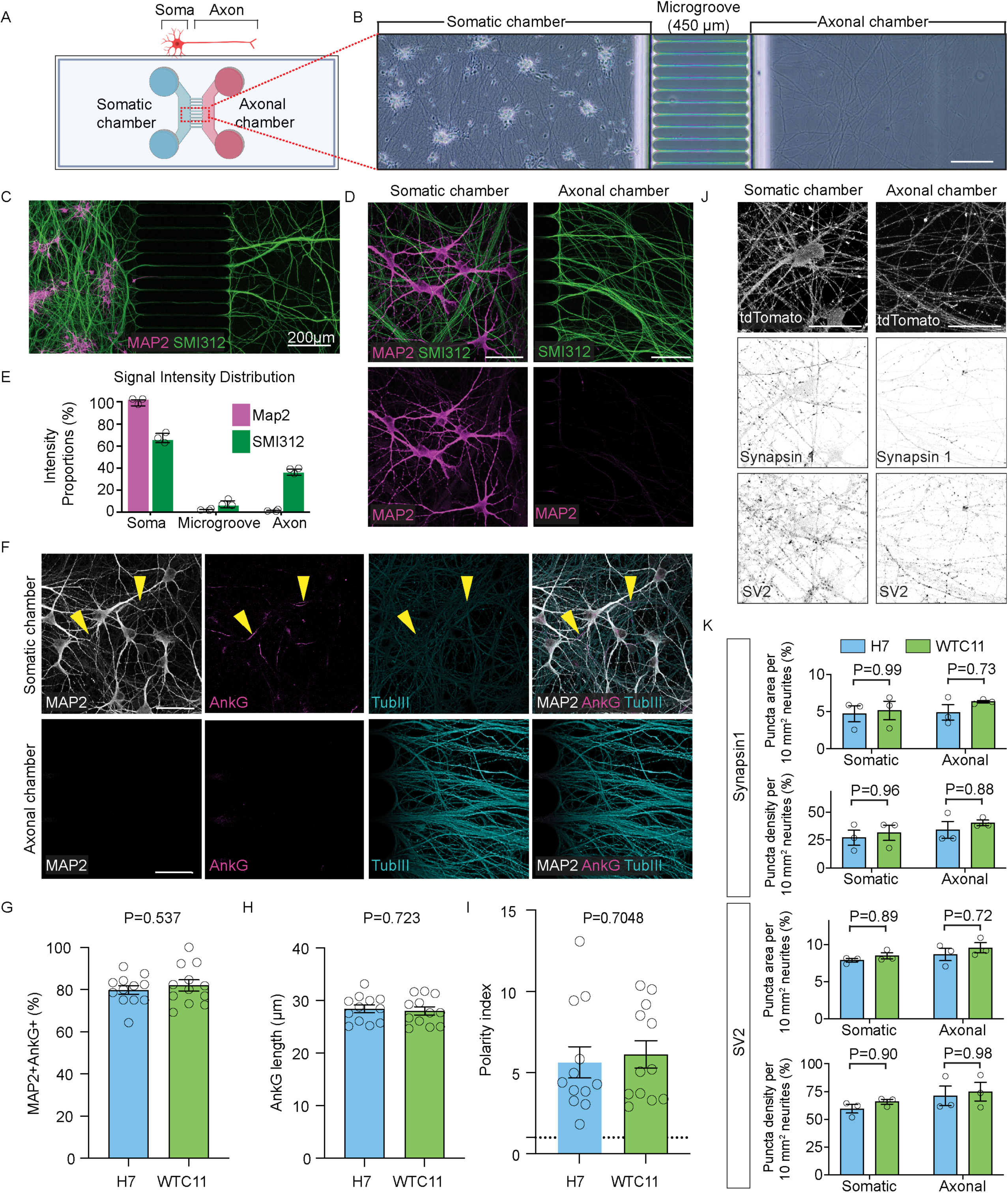
hRGCs show proper polarization in a microfluidics system. A-B. Schematic. (**A**) and representative bright field images (**B**) of the organization of the microfluidics system. hRGCs are plated in the somatic chamber and extend axons through 450 um microgroove to the axonal chamber. Scale bars = 200µm. **C-D**. Representative low magnification (**C**) and high magnification (**D**) images of hRGCs grown in the microfluidics system with the dendritic marker MAP2 and the axonal marker SMI312 labeling. Scale bars = 50µm. **E**. Quantification of MAP2 and SMI312 intensity distribution in microfluidics device**. F.** Representative images of axon maturation show that hRGCs develop defined axon initial segments that are proximal to the cell body, localized to axons [TubIII positive (cyan) while MAP2 negative (greyscale)] and positive for structural protein AnkG (magenta, arrows). Scale bars = 50µm. **G-I**. Quantification of hRGCs in microfluidics chambers derived from both hPSC lines show that most cells possess an AIS (**G**) and that the length of the AIS is approximately ∼30um (**H)**. There is no significant difference in these values between hRGCs derived from the two H7 and WTC11 hPSC lines. **I.** The polarity index was calculated as the intensity ratio of MAP2 and AnkG. Both cell lines exhibited strong polarization, with polarity index values ranging from ∼1,5 to 3.5, and no significant differences were observed between the hRGC groups (n = 12 images from 3 biological replications, each image contains >10 hRGCs). **J-K**. Representative images (**J**) and quantification (**K**) of the distribution, puncta area and puncta density of the synapse vesicle protein SV2 and synapse protein SYN1 in the somatic versus the axonal chamber of hRGCs cultured in the microfluidics system. Scale bars = 50µm. No significant differences were observed between the two lines or in the two compartments, though the density of SV2 positive puncta tended to be higher than that of SYN1 (n = 3 biological replications, each replicate was quantified from 4 images). Data are presented as average values ± the s.e.m.

As in the standard culture system, hRGCs in the microfluidics device developed highly ordered AISs positive for the AIS marker AnkG, which was present in axons, but not in dendrites, as indicated by colocalization with Tubulin III (**Figure 2F**). Quantification revealed that similar to standard hRGC cultures, ∼80 % of hRGCs contained an AIS and no significant differences were found between the two lines (*p = 0.537,* **Figure 2G**). The length of the AIS in microfluidics hRGCs mirrored that in the standard culture system, at 28 ± 1.06 µm, though in this case lengths did not vary between hRGCs derived from the two cell lines (*p = 0.723,* **Figure 2H**). hRGCs grown in the microfluidics system were also positive for the AIS enriched proteins β-spectrin and sodium channels (**Supplemental Figure 2B**). Analysis of the polarity index in these cells revealed that hRGC axonal segregation in the microfluidics system was also robust, with values from ∼1.5-∼13, and no significant difference was observed between the cell lines (*p = 0.7048,* **Figure 2I**).

We next asked whether hRGCs had a baseline potential to form synapse proteins by staining for Syn1 and the synaptic vesicle protein SV2. Both proteins could be detected in the somatic and axonal compartments and formed punctate localizations that overlapped with neurites in both chambers (**Figure 2J**). Quantification of the puncta area and puncta density for both markers showed that area and density of SV2 was higher on average than that of Syn1. In both cases, no significant difference was observed between the somatic and the axonal chamber nor were there notable variations in values between the two hPSC lines (**Figure 2K**). Together, these data show that hRGCs from both hPSC lines can be induced to spatially segregate their dendrites and axons, modeling the arrangement that occurs *in vivo*, and show proper polarization and AIS formation in these systems.

### Retinorecipient targets promote region-specific hRGC connectivity that models *in vivo* innervation

During visual system development, RGCs extend axons that specifically and differentially connect with retinorecipient areas and avoid connecting with non-visual targets. In both mice and humans, most RGCs form connections in the LGN, while the SCN is innervated by fewer RGC types^3^. We thus asked whether these distinctions in connectivity could be preserved in an *in vitro* eye-brain connectivity model. Given the lack of human iPSC derived region-specific visual targets^33,34^, we bioinformatically evaluated the suitability of mouse visual targets as surrogates relative to available thalamic organoid models. Specifically, we computed, the reference similarity spectrum^35,36^ using publicly available single-cell and single-nucleus RNA-seq data of neuronal populations from developing mouse LGN and human iPSC-derived thalamic organoids, relative to primary LGN transcriptomic profiles found in the Human Brain Cell Atlas. We found that the mean score for the developing mouse dLGN (0.63) was comparable to that of one thalamic organoid dataset (0.64), though the organoid data required pre-filtering to remove non-specific cells generated during differentiation (**Figure 3A**). In contrast, mouse dLGN scored notably higher than the thalamus region from the Human Neural Organoid Cell Atlas (0.41). Furthermore, the high-performing organoid dataset exhibited a substantially wider distribution of similarity scores compared to the mouse dLGN (**Figure 3B**). This broad distribution reflects heterogeneity in the organoid model, whereas the tighter distribution of the mouse dLGN indicates a more homogeneous population of relevant postsynaptic neurons.

**Figure 3.**
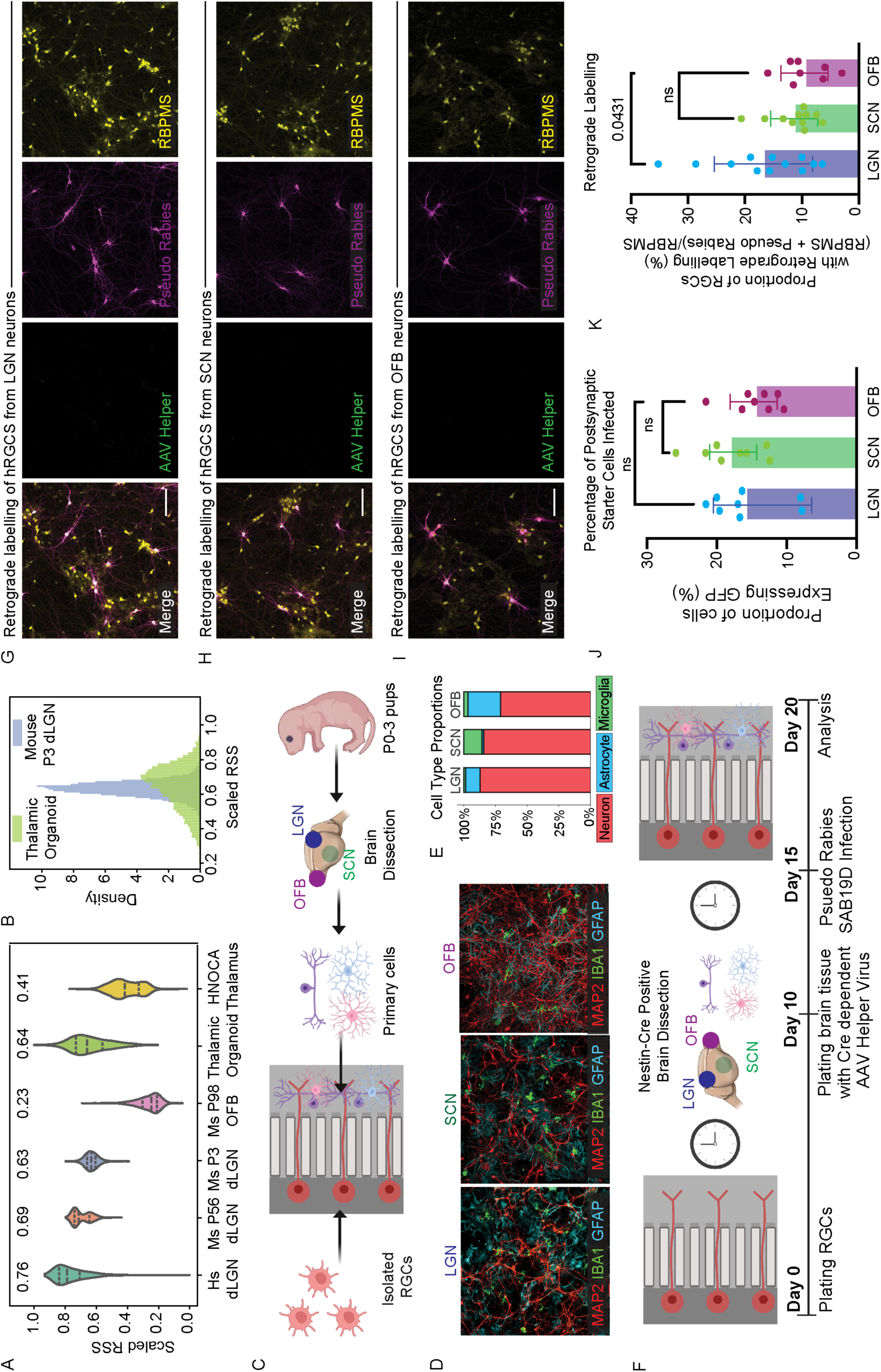
hRGCs form brain region specific connections with post-synaptic targets. **A**. Relative similarity score of how well mouse brains and thalamic organoids represent human single-cell data. **B**. Density distribution map illustrates variability across publicly available datasets. **C.** Schematic of the generation and addition of retinorecipient and control brain regions to the microfluidics system. **D-E.** Major cell type representation in the retinorecipient and control brain regions. Cultures from each region [lateral geniculate nucleus (LGN), suprachiasmatic nucleus (SCN) and olfactory bulb (OFB)] were comprised primarily of neurons (MAP2, red, ∼75-85%), while microglia (Iba, green), and astrocytes (GFAP, cyan) were also present. n = 8 images from 3 biological replicates. Scale bars = 50 µm **F**. Schematic of retrograde circuit tracing approach. hRGCs were grown in microfluidics chambers for 10 days. Following this period, dissociated cells from dissected control brain regions (OFB) and retinorecipient brain regions (LGN and SCN) were isolated from postnatal day (P) 0 to P3 mouse pups, infected with AAV-helper GFP,, and then plated in the axonal chamber. 5 days later, dissected brain regions were infected with the pseudo rabies virus. **G-I.** Representative images of the number of transsynaptic labeled following retrograde tracing from retinorecipient LGN and SCN neurons or OFB control neurons. Scale bars = 50 µm. **J**. The number of starter brain cells that were infected with the Cre dependent AAV helper virus were not statistically significant across different starter brain cells. **K.** While a degree of connectivity was observed in all cases, LGN neurons tended to have higher degrees of connectivity relative to control OFB neurons (P=0.0431).

Based on these results, we chose to isolate and utilize the developing LGN and SCN from neonatal (postnatal day, P) P1-P3 mice (**Figure 3C**). As a non-retinorecipient control brain region, we chose to isolate the olfactory bulb (OFB). To ensure isolated brain regions were specific to the LGN and SCN, we first performed anterograde tracings using cholera toxin B (CTB) injected in the eyes of P3 pups. We found labeled RGC axons populated isolated LGN and SCN but not the OFB, validating brain target identity (**Supplemental Figure 3**). We then confirmed that cultured mouse LGN, SCN and OFB retained the normal distribution of cell types and area-specific neuron subtype markers observed in these regions using immunohistochemistry (**Figure 3D-E**). The bulk of cells in these cultures were neurons, as indicated by staining for MAP2 (∼70-85%), and microglia (IBA1) and astrocytes (GFAP) were also present at expected ratios (**Figure 3D-E**)^37^. In addition, a subset of cultured LGN neurons were positive for FOXP2, a transcription factor enriched in a some LGN neurons *in vivo*^38,39^, while a subset of SCN cells were positive for vasoactive intestinal polypeptide (VIP), neurons essential for circadian entrainment^40,41^ **(Supplemental Figure 4A-B**). These data are consistent with the idea that post-synaptic cellular identity is preserved in LGN and SCN cultures.

We next asked whether hRGCs could form connections with postsynaptic targets. To assess this, we focused on H7-derived hRGGs as there were few significant differences between hRGCs from both lines, and the H7 line had the advantage of carrying the tdTomato reporter. H7-derived hRGCs were plated in the somatic chamber of the microfluidics system and allowed to extend axons, followed by the addition of LGN, SCN, or OFB cells into the axonal chamber. We found that in all cases, tdTomato hRGC axons anatomically colocalized with MAP2 positive postsynaptic target neurites, suggesting the potential for connectivity (**Supplemental Figure 5**). We then assessed connectivity directly using retrograde viral tracing with the Rabies/SAB19D system (**Figure 3F-K**). In this approach, we leveraged Cre containing post-synaptic neurons (LGN, SCN, or OFB) derived from Nestin-Cre mice. Following isolation, we infected these ‘starter neurons’ with a Cre-dependent AAV helper virus containing the TVA receptor for rabies virus entry and the rabies glycoprotein for spread. These cells were then plated in the axonal chamber of microfluidics devices containing WTC11 hRGCs. Five days later, Rabies/SAB19D containing an mCherry tag was added to the postsynaptic neurons. No significant differences in infection rates were observed in starter neuron populations across brain target regions (**Supplemental Figure 6, Figure 3J**. Retrograde spread was assessed in RGCs in the somatic compartment after an additional 5-day incubation period. This system ensures specificity and directionality, as only Cre containing post-synaptic cells will serve as starter neurons through which rabies can spread retrogradely through synapses to Cre negative presynaptic cells (**Figure 3F-K**). We found that the number of hRGC somas that were labeled by retrograde tracing was significantly elevated when LGN targets were provided relative to SCN or OFB (**Figure 3K**, P = 0.0431). These results were supported by parallel experiments using a AAV-Retro GFP virus, which showed that the number of hRGC somas that were labeled by retrograde tracing was elevated in presence of LGN and SCN postsynaptic targets relative to OFB (**Supplemental Figure 7**). These data suggest that presynaptic hRGCs axons can form properly orientated connections and that hRGCs preferentially connect with retinorecipient neurons in a microfluidics model of eye-brain wiring.

To validate and extend these findings, we asked whether the area and number of synapses formed between hRGCs differed between LGN, SCN, and OFB. As above, hRGCs were grown in microfluidics chambers and allowed to extend axons, followed by the addition of LGN, SCN or OFB targets (**Figure 4A**). Ten days after post-synaptic target addition, Syn1 was used to label and quantify the area and number of synapses formed on tdTomato positive RGC axons.

**Figure 4:**
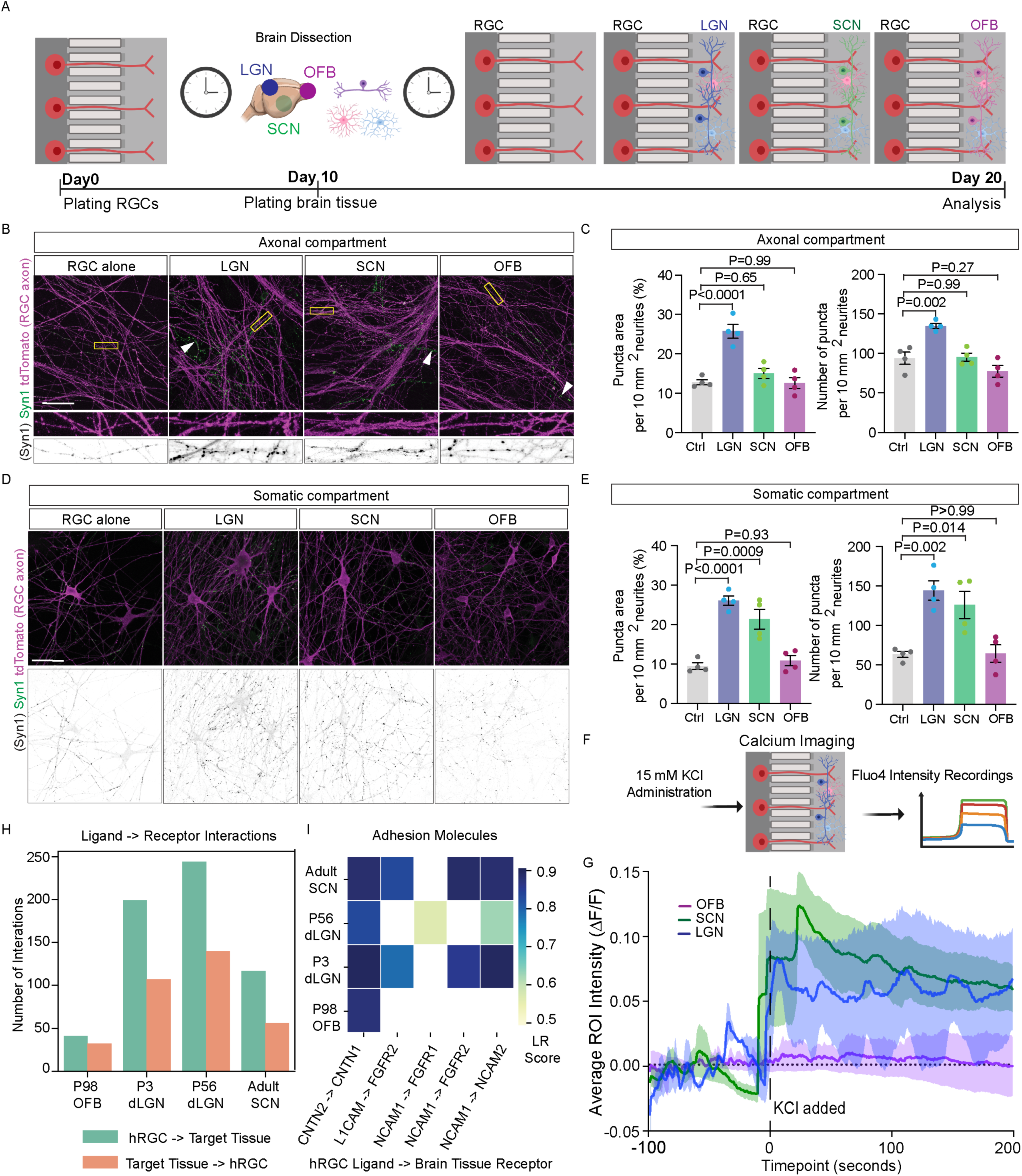
Retinorecipient targets specifically promote hRGC synapse formation. **A**. Schematic of the microfluidics system following the addition of diverse post-synaptic targets to the axonal chamber, lateral geniculate nucleus (LGN), suprachiasmatic nucleus (SCN) and olfactory bulb (OFB). **B-C**. Representative images (**B**) and quantification (**C**) of Synapsin 1 (SYN1, green) along hRGC axons (tdTomato, magenta) following the addition of dissociated cells from retinorecipient and control brain regions to the axonal chamber (white arrows). The addition of LGN partners significantly enhanced Syn1 puncta area and puncta number relative to hRGCs alone (Ctrl), SCN, or OFB partners (n = 4 biological replications, each replicate was quantified from 4 images). Scale bars = 50 µm. **D-E**. Representative images (**D**) and quantification (**E**) of Synapsin 1 (Syn1, GFP) along hRGC dendrites (tdTomato, magenta) in the somatic chamber following the addition of dissociated cells from retinorecipient and control brain regions to the axonal chamber. The addition of either retinorecipient partner (LGN, SCN) significantly enhanced hRGC maturation as indicated by the increase in Syn1 puncta area and puncta number relative to hRGCs alone (Ctrl) or OFB partners.n = 4 biological replicates, and 4 images were used to quantify each replicate. Scale bars = 50 µm. Data are presented as average values ± the s.e.m. **F-G**. Schematic of calcium imaging experiment (**F**) and Calcium imaging intensity signalling [NT1] of OFB, SCN, and LGN brain cells connected to hRGCs (**G**). 15 mM KCl was used to synchronously depolarize hRGCs in the somal compartment and measure calcium changes in the postsynaptic neurons. Both LGN and SCN postsynaptic neurons produced a robust response to hRGC depolarization, while OFB postsynaptic neurons showed little to no response. **H-I**. Ligand-Receptor analysis shows that there are significantly more ligand-receptor interactions between hRGCs and SCN or LGN brain regions than OFB brain regions (**H**) and that canonical adhesion molecules have been identified (**I**).

We found that the addition of LGN neurons significantly increased the number and size of Syn1 positive puncta in the axonal compartment relative to hRGCs alone or the addition of SCN or OFB cells (P< 0.0001; **Figure 4B-C).** We also assessed Syn1 puncta number and area in the somatic compartment as one read out of the impact of postsynaptic neurons on presynaptic neuron maturation. We found that the addition of either LGN or SCN neurons significantly increased the neurite area occupied by Syn1 (P *≤* 0.001) and the absolute number of Syn1 positive punction (P *≤* 0.01) relative to either hRGCs alone or the addition of OFB cells (**Figure 4D-E**).

To directly assess functional connectivity, we then undertook cytosolic calcium imaging experiments in our microfluidics systems. Ten days after the addition of LGN, SCN, or OFB to RGCs axons, we loaded post-synaptic neurons in the axonal chamber with the calcium indicator Fluo-4. We then used KCl (15 mM final concentration) to synchronously depolarize hRGCs in the somatic compartment and measured calcium changes in the post-synaptic neurons in the axonal compartment (**Figure 4F)**. We found that hRGC depolarization resulted in robust calcium responses in both LGN and SCN target neurons, but not in OFB neurons (**Figure 4G).** Together these experiments show that hRGCs can fire action potentials that drive neurotransmitter release from presynaptic axonal terminals which cause calcium influx in retinorecipient but not control target neurons. Further, when LGN, SCN, and OFB neurons were exposed to hRGC axons, post-synaptic neuron maturation improved as measured by the number of postsynaptic neurons that contained AnkG positive axon initial segments (**Supplemental Figure 8**). RGC axons also appeared to promote AIS shortening across post-synaptic targets, consistent with a known correlation between AIS length and neuron maturation^42–45^. Together with our connectivity and functional analyses described above, these results support the idea that RGC axons can promote the maturation of co-cultured neurons across different brain regions but that RGCs structurally and functionally connect specifically with retinorecipient regions. To investigate possible biological mechanisms underlying the specificity of this synaptic formation, we performed cell-cell communication analysis using publicly available transcriptomic data^46,47^. We assessed potential interactions between hRGC datasets and mouse LGN, SCN, and OFB datasets using LIANA+^46^. We found that number of predicted ligand-receptor interactions was higher between hRGCs and either LGN or SCN than between hRGCs and OFB (**Figure 4H**). This target specificity was maintained regardless of directionality (hRGCs defined as the ligand-producing source or the receptor-expressing target), supporting the existence of robust, bidirectional signaling between the retina and visual brain. Furthermore, our analysis enriched for canonical cell adhesion interactions known to mediate circuit assembly, including CNTN2–CNTN1, L1CAM–FGFR2, NCAM1–FGFR1, and NCAM1–NCAM2^48–50^ (**Figure 4I**). Collectively, these transcriptomic findings, paired with our circuit tracing and functional assays, confirm that the molecular logic of *in vivo* visual connectivity is preserved in our *in vitro* system, validating its utility for future discovery.

## Discussion

To understand human neural development and disease so that effective therapies can be developed, it is important to have robust models of neural connectivity that preserve cell identity, wiring patterns, circuit organization, and specificity. In this study, we developed an *in vitro* microfluidics model of eye-to-brain connectivity using hPSC-derived RGCs to evaluate connectivity characteristics specific to different postsynaptic brain regions. To understand inherit variability of different stem cell lines, we used two different hPSC lines to show that they can give rise to hRGCs that develop polarized dendritic and axonal structures when cultured alone and when plated in microfluidics systems. Axons grew long distances (>450 um), effectively targeted the distal chamber, and formed distinct axon initial segments that were of similar lengths and organization to that observed in human samples *in vivo*. To assess the specificity of hRGC connectivity with particular brain visual targets, we chose to use developing LGN and SCN from neonatal mice given their transcriptional similarity to parallel human targets the lack of human iPSC derived region-specific visual targets^33,34^. We found that hRGCs can differentially sense and respond to mouse retinorecipient targets to control wiring outcomes. Retrograde rabies virus tracing showed enhanced connectivity from retinorecipient targets relative to control regions, and synapse number and puncta size were also elevated when LGN and SCN were provided. Furthermore, functional activity appears to be restricted to retinorecipient targets only, as indicated by calcium response measures in our system. These findings indicate that the innervation and specificity properties of human RGCs are maintained in our culture systems, allowing these cells to selectively recognize and respond to retinorecipient targets. These data are thus consistent with an instructive model of connectivity. This and related systems will thus be valuable in identifying human-specific wiring factors for potential therapeutic interventions.

There are several advantages to our model of eye-to-brain connectivity. First, it has the capacity to enable structurally separated pre- and postsynaptic manipulation and assessments of human RGCs for both normal development and in the context of genetic disease variants. This is an important advancement because while other studies have shown that retinal organoids can give rise to RGCs, these cells die at later stages of differentiation^15,16^. Further, hRGC axons do not develop well in organoids and are unable exit organoids under standard preparations, precluding the ability to study axon initiation, extension and connectivity^15,16^. While several factors likely contribute to hRGC loss in these systems, one factor may be the lack of neurotrophic support from postsynaptic partners^51^. Our model overcomes these limitations by providing hRGCs with appropriate postsynaptic partners. A second advantage of our system is the use of distinct retinorecipient targets derived from mice that preserve region-specific neural populations, allowing for the examination of differential innervation properties. This approach is particularly valuable, as current human thalamic culture systems do not facilitate the derivation of LGN versus SCN cell types nor do they include glia, such as astrocytes and microglia^52^, which are known to impact visual circuit connectivity^53,54^. Third, we can precisely control the developmental timing of post-synaptic partners to model *in vivo* events. In mouse, RGC axon terminal synapses are formed and refined with their postsynaptic partners from ∼E16-P12, which overlaps with the P0-P3 timepoint from which we harvest post-synaptic target cells ^55,56^. Fourth, we can take advantage of the numerous genetically modified mouse models to generate target neurons with various modifications. For example, here we utilize Nestin-Cre mice to enable selective, Cre-dependent retrograde tracing from post-synaptic neurons to hRGCs. This facilitates molecular and functional studies. Finally, our microfluidics system permits direct assessment of directional connectivity, synapse number, and synapse size, which can prove challenging in other approaches. This model provides an alternative to human assembloid circuits^16^ and related systems^57–59^ where directional connectivity cannot be precicely controlled.

How might human eye-brain connectivity models be used to understand RGC axonal guidance and postsynaptic interactions? In the developing eye, RGC axons exit the eye, form the optic nerve, determine whether to project ipsilaterally or contralaterally at the optic chiasm, and form connections with specific brain targets. Several repulsive and attractive cues help guide this process, including Eph receptors and ephrins (reviewed in^60^). In addition, the adult SCN expresses slit/robo inhibitory cues that prevent innervation from RGCs^61^. Mining publicly available single-cell RNA Sequencing datasets from stem cell derived hRGCs and retina-specific brain targets, we showed ligand-receptor pairings that are canonical and align with molecular cues found in literature. Based on these results, it is interesting to envision alternative forms of the system we describe that could be used to model various aspects of axon guidance and connectivity. These include: 1) providing cues in alternative chambers, 2) allowing axons to ‘choose’ between different retinorecipient targets, or 3) modeling competitive axon refinement between ‘ipsilateral’ and ‘contralateral’ RGC inputs. In addition, this system may be suitable for uncovering brain region specific factors that provide RGC trophic support. For example, our data are consistent with the idea that postsynaptic partners enhance RGC maturation as the addition of retinorecipent targets but not a non-recipient control brain region improved of Syn1 positivity in the RGC somatic compartment. In line with this idea, retrograde neurotrophic support from postsynaptic neurons can also directly enhance RGC survival both in development^62–64^ and in disease conditions^65,66^. Lastly, the capability to examine distinct post-synaptic targets has the added advantage of providing a system to study RGC influence on brain neuron maturation. For example, RGC axons can provide a scaffold for LGN maturation via activity dependent regulation of astrocytes and their ability to produce the inhibitory neuron recruiting molecule fibroblast growth factor 15^67^. In future studies, it will be interesting to assess the long-term effects of individual postsynaptic targets in modulating RGC persistence and resilience, and conversely to assess the impact of RGCs on brain neuron maturation.

Finally, the system we describe has potential utility for the study of retinal and other neurodegenerative diseases. For example, human patient samples could be used to grow hRGCs to model individual diseases and phenotypes^68,69^. This is particularly important because the expression of proteins linked to genetic glaucoma risk in humans (e.g. such as *OPTN*^70^*, GAS7*^71^*, SIX6*^72^, and *MAP3K1*^73^) may not be well conserved in mouse RGCs. This is supported by findings that genetic models of human glaucoma risk factors, like the E50K mutation in OPTN, result in only mild phenotypic changes in mice^74^, despite causing severe disease in humans^75^. In addition, our system permits compartmentalized study and manipulation of dendritic versus axonal hRGC outcomes. This is of relevance to diseases such as glaucoma, in which both RGC dendrites and axons degenerate, and these processes may occur via different mechanisms^6,7^. Another important application will be the discovery of human molecular pathways that determine brain-region specific connectivity outcomes and resiliency. The potential for this system as a discovery platform is particularly encouraging given that RGC axons formed a significantly higher number of Syn1 positive puncta with the LGN compared to the SCN, modeling the higher number of RGCs that connect with the LGN versus the SCN in primates^17,76^. It will also be interesting to determine whether RGCs that innervate LGN versus SCN postsynaptic targets are differently resilient to disease-related insults, as they are *in vivo*^77^. Such studies may provide new therapeutic opportunities for precise intervention in the progression of retinal and other neurological diseases.

## Materials and Methods

### hPSCs maintenance and retinal organoid differentiation

Two hPSC lines, human embryonic stem cell line H7 and human induced pluripotent stem cell line WTC11, were used in this manuscript. H7 has been previously genetically engineered with an RGC tdTomato reporter and Thy1.2 antigen at the BRN3b locus as described (BRN3:tdTomato:Thy1.2)^19^, while WTC11 was edited using similar approaches to genetically introduce the Thy1.2 antigen without tdTomato at the BRN3b locus (BRN3:Thy1.2). hPSCs were maintained in Matrigel-(Fisher Scientific, cat# 8774552) coated plates with mTeSR™ Plus (StemCell Technologies, cat# 100-0276) medium. hPSCs were split every 4-5 days by Dispase (2 mg/mL, Life Technologies, cat# 17105041) as described previously^69,78–80^.

Methods for generating retinal organoids have been described previously ^69,78–80^. In brief, to generate retinal organoids, hPSCs at 80% confluence were enzymatically dissociated with Dispase for 10 min in a 37°C incubator. Dissociated hPSC clumps were plated on low adherent plates in mTeSR1 (StemCell Technologies, cat# 85850) and Neuronal Induction Medium [NIM, composed of DMEM/F12(1:1) supplemented with 1X N2, MEM-NEAA, and 2 µg/ml heparin] with a ratio of 3:1 on day 0 and gradually transitioned to full NIM on day 3 (mTeSR1 to NIM at 1:1 on day 1 and at 1:3 on day 2) to induce embryonic bodies. On day 6 the medium was changed to NIM supplemented with BMP4 (50 ng/mL, R&D Systems, cat# 314-BP-050) to enrich retinal organoids. Embryonic bodies were plated on 6-well plates on day 8 at a density of 100-200 embryonic bodies per well in NIM with 10% FBS to promote cell adherence. NIM was replaced the next day, and the medium was then changed every 2-3 days. On day 16, the loosely attached rosettes were mechanically lifted using a P1000 tip, and colonies were transferred to low-adherent plates to allow retinal organoid specification. Colonies were fed with fresh Retinal Differentiation Medium [RDM; composed of DMEM/F12 supplemented with 1X B27, 1X MEM-NEAA, and 1X Antibiotic-Antimycotic (ThermoFisher, cat#15240062)] every 2-3 days until the organoids were collected. RDM was supplemented with 1X GlutaMAX starting at day 20. FBS was also supplemented to the RDM, but in incremental concentrations, with 1% FBS on day 20, 5% FBS on day 25, and 10% FBS on day 30. After day 35, retinal organoids were fed every 2-3 days with RDM containing 10% FBS, 1X GlutaMAX, and 100 μM Taurine.

### RGC purification from retinal organoids

Methods for RGC isolation and purification from retinal organoids have been described previously with slight modifications^19,69^. Between days 35 and 60, retinal organoids were collected in a 15 mL conical tube, washed with warm DMEM/F12, and incubated with dissociation solution in a water bath at 37°C for 20 min. Cells were agitated every 5 min. The dissociation solution was composed of AccuMAX (Life Technologies, cat# 00-4666-56), Papain (16 unit/mL, Worthington Biochemical Corporation, cat# LS003126), DNase I (Sigma, cat# D4527), 5.5mM L-cysteine (Sigma, cat# C1276), and 100X HEPES (Life Technologies, cat# 15630080). The dissociation solution was activated by incubation in a water bath at 37°C for 15 min prior to introduction of organoids. After incubation with retinal organoids, dissociation solution was replaced with RDM supplemented with 10 mg ovomucoid inhibitor (Worthington Biochemical Corporation, cat# LS003087) and 10 mg BSA. Retinal organoids were mechanically dissociated with a P1000 pipette, and single cells were strained through a 30 µm filter (Miltenyi, cat# 130-041-407). After collection, dissociated filtered cells were centrifuged and washed with RDM. Using a MACS cell separation kit^19^, cells were incubated with CD90.2 microbeads (Miltenyi, cat# 130-121-278) for 20 minutes before being passed through MACS magnetic bead purification columns. After purifying cells, 30-50k RGCs were plated on a poly-D-Lysine (Life Technologies, cat# A3890401) and Laminin (Life Technologies, cat# 23017015)-coated 12 mm coverslip in Neurobasal-A medium (Life Technologies, cat# 10888022), supplemented with 1X N2, B27, GlutaMAX, 1X Antibiotic-Antimycotic, 20 ng/mL BDNF, and 20 ng/mL GDNF. The medium was changed every 3-4 days and cells were maintained for up to 4 weeks.

### RGC-brain postsynaptic target cultures in microfluidics devices

Microfluidics devices (Xona Microfluidics, cat# XC450) were set up following manual instructions. On day 1, microfluidics devices were treated with XC Pre-Coat solution and coated with poly-D-lysine overnight at 4°C. On day 2, devices were washed with PBS three times and coated with Laminin overnight in a 37°C incubator. The following day, 50-100k purified RGCs were plated on the device. To maintain the fluidic physical isolation, somatic chambers were each given 150 µl of Neurobasal-A medium, while axonal chambers were each given 100 µl of Neurobasal-A medium. To induce axon outgrowth, 50 ng/mL BDNF and 50 ng/mL CNTF were supplemented in the axonal chamber^16^. The medium was changed every 2-3 days and maintained for up to 4 weeks. For RGC-brain postsynaptic target cultures, 80k RGCs were plated in the somatic chamber for an initial 10-day period. This was followed by plating ∼100K of dissociated brain target cells in the axonal chambers for an additional 10 days before the cultures were collected for analysis. Brain target isolation and dissociation are described below.

### Animal models and primary cultures

Experiments with mice were carried out in accordance with the recommendations in the Guide for the Care and Use of Laboratory Animals of the NIH under protocols approved by the BCM Institutional Animal Care and Use Committee. C57BL/6J strain (JAX 000664) or CD1 wildtype mice (obtained from Charles River Laboratories) were used in this study. To establish the primary cultures, brain tissue from P0-P3 mice were dissected from LGN, SCN, and OFB brain regions and collected in cold DMEM/F12 medium. After dissection, tissues were incubated with papain solution (DMEM/F12 medium containing 16 unit/mL papain, DNase I, L-cysteine, and 100X HEPES) in a water bath at 37°C for 20 min. Cells were agitated every 5 min. After incubation was complete, papain solution was replaced and quenched by DMEM/F12 supplemented with 10 mg ovomucoid inhibitor and 10 mg BSA. Brain tissues were mechanically dissociated with a P1000 pipette, and single cells were strained through a 70 µm filter. 50k cells were then plated in poly-d-lysine and laminin-coated 12 mm coverslips for cultures of brain targets alone, while 100k cells were plated in the axonal chamber of microfluidics devices with Neurobasal-A medium. Media was changed every 3-4 days and maintained for up to 10 days.

### Anterograde Labeling Eye Injections

For anterograde brain target labelling experiments, cholera Toxin Subunit B (CTB), Alexa Flour 488 conjugate (cat# C34775) was resuspended to a final concentration of 1 ug/uL in PBS. P0-P3 aged mice were injected with 1 uL of CTB solution and allowed to recover. After 24 hours, pups were collected, and the LGN, SCN, and OFB brain regions were dissected out and placed directly on a slide. Collected samples were then imaged at 10X magnification using the Olympus Fluoview FV1200 confocal microscope.

### Viral transsynaptic tracing

For retrograde Rabies virus tracing, approximately 100,000 hRGCs were seeded and maintained in the microfluidics system for 10 days. At the 10-day timepoint, ∼100,000 cells were isolated and dissociated from the SCN, LGN, and OFB (control) of Nestin-Cre positive animals at P0-P3 and seeded onto the axonal compartment of microfluidics devices. For each batch of seeded cells, we then applied 10 uL of conditional AAV-FLEX-GTB virus (Final concentration: 2.85E+9 GC/mL, packaged in AAV2) to allow for Cre-dependent TVA, Rabies G protein, and GFP expression. After 5 days of initial AAV infection, we then performed a secondary infection with 10uL of mCherry-tagged pseudotyped Rabies Virus (EnvA G-Deleted Rabies-mCherry, final concentration: 2.00 E+10 GC/mL, packaged at Salk Institute)^81,82^ to allow retrograde tracing of monosynaptic inputs to the seeded populations of cells. We then assessed the number of mCherry reporter postivie hRGC somas as a measure of monosynaptic connectivity between hRGCs and their post-synaptic targets.

AAV viruses were packaged with pAAV-Syn1>EGFP:WPRE (VectorBuilder, cat# VB240311-1387gfy). For retrograde tracing, brain target cells (LGN, SCN, or OFB) were infected with AAV2-retro-Syn1>EGFP:WPRE at an MOI of 1 x 10^6^. All the AAV viruses were packaged by VectorBuilder. Cells were infected in 96-well U-bottom plates and incubated in a 37°C incubator for one hour. After viral infection, the cells were washed with warm DMEM/F12 three times to remove any residual AAV from the medium. Cells were resuspended in Neurobasal-A medium and plated in the axonal compartment of microfluidics devices. To ensure the proper viral transsynaptic tracing and no residual AAV in the culture medium, the medium from the last wash was collected and added to microfluidics devices as a negative control.

### Calcium Imaging

For calcium imaging, approximately 100,000 hRGCs were seeded and maintained in the microfluidics system for 10 days. At the 10-day timepoint, ∼100,000 cells were isolated and dissociated from the SCN, LGN, and OFB (control) pups at P0-P3 and seeded onto the axonal compartment of microfluidics devices, where they were maintained for an additional 10 days. For calcium imaging, cells were first washed with physiological saline containing: 140 mM NaCl, 5.6 mM KCl, 1.6 mM CaCl2, 1 mM MgCl2, 10 mM glucose, and 10 mM HEPES, pH 7.3. Next, Fluo-4 acetoxymethyl ester (Fluo-4 AM; Thermofisher cat# F14201) was added to the axonal chambers at a final concentration of 3 uM. Cells were then incubated in the dark at room temperature for one hour to allow for adequate loading. After one hour, cells were washed with the same solution and incubated for an additional 30 minutes before data acquisition began. For data acquisition, 15 mM KCl was used to synchronously depolarize hRGCs in the somal compartment and measure calcium changes in the postsynaptic neurons. The Fluo-4 loaded cells were identified and imaged using Slice Scope Pro 6000 (Scientifica, UK) platform, equipped with optiMOS camera (QImaging) controlled by Micro-Manager version 1.4.22 software (University of California at San Francisco). Thorlabs DC4100 470-nm LED illumination setup with 49002-ET-EGFP (FITC/Cy2) emission filters (Chroma Technology) were used as a source of excitation light. The resulting fluorescent images were acquired in time lapse acquisition mode at 1 image per second. The resulting image tack files were analyzed offline using FIJI by measuring the average calcium intensity changes over time. Each condition was performed in triplicates, and 30 ROIs were use for each image tack file.

### Immunocytochemistry

For staining axon initial segment proteins, the cells were fixed with 2% paraformaldehyde at room temperature for 15 min. For all other staining, cells were fixed with 4% paraformaldehyde at room temperature for 30 min, followed by blocking with 10% donkey serum, 5% BSA, and 0.2% Triton-X in PBS for an hour. Primary antibodies were diluted in 5% donkey serum, 2.5% BSA, 0.1% Triton-X, and PBS and incubated overnight at 4°C. The following day, cells were washed with PBS three times before being incubated at room temperature for 1 hour with secondary staining solution [secondary antibodies and DAPI (Cell Signaling, cat# 4083, 1:1000) diluted in in 5% donkey serum, 2.5% BSA, 0.1% Triton-X, and PBS]. The cells were then washed with PBS three times and stored either in mounting media (ProLong Gold Antifade Mountant, Life Technologies, cat# P36930) or PBS (for microfluidics devices). Images were acquired using an Olympus Fluoview FV1200 confocal microscope, ZEISS LSM 880 confocal microscope, or ZEISS Axio Observer. All images were processed in Fiji (ImageJ).

The following antibodies were used in our analyses:

### Polarity index quantification

The AIS was defined by AnkG labeling, while dendrites were defined by MAP2 labeling. To calculate the polarity index, we used the AnkG signal in the axon of an individual hRGC to create a traced object. This traced object was then applied to 3 proximal dendrites of the same neuron (Map2 positive neurites). From these values, the polarity index was calculated by dividing the AnkG intensity in the axon by the average AnkG intensity in the dendrites. This metric was then performed on 8-10 neurons per condition to arrive at the average polarity index per condition. Ratios greater than 1 indicate that AnkG expression is polarized in the axon. AnkG expression along a neurite was traced to create the region of interest. In the region of interest, the intensity of AnkG was measured using Fiji-ImageJ software. The polarity index was calculated as the ratio of AnkG intensity in an axon to AnkG intensity in a dendrite. At least 9 neurons were measured for each condition.

### Synapse quantification

Synaptic density quantification was performed in Fiji. tdTomato fluorescence was used to visualize the neurites and determine the neurite area, while Syn1 fluorescence was used visualize synaptic puncta. Puncta number and area were quantified using Fiji particle analyzer. To quantify hRGCs axonal Syn1 levels in microfluidics devices, a color threshold of merged tdTomato and Syn1 images was set before using the particle analyzer. This step enabled quantification of Syn1 levels specifically in hRGC axons and excluded Syn1 in LGN, SCN, or OFB neurons.

### Neurite reconstruction

hRGCs in 12mm coverslips were transfected by Lipofectamine 3000 (Life Technologies, cat# L3000001) with 500 µg of pAAV-hSyn1-EGFP-P2A-EGFPf-WPRE-HGHpA plasmid (Addgene, cat# 74513) following manual instructions. Fresh medium was replaced after 6 hours of transfection to prevent cell toxicity. hRGCs were fixed after 48 hours of transfection following immunocytochemistry with GFP antibody staining. Images were acquired using ZEISS Axio Observer and processed in Fiji using a neuroanatomy plug-in for neurite tracing and Sholl analysis.

### Statistical analysis

All experimental data are presented as mean ± s.e.m., with n indicating the number of replicates across all experiments. Statistical analyses were performed using either Student’s t-test or ANOVA followed by Tukey’s post hoc test, utilizing GraphPad Prism 10. Statistical significance was defined as p < 0.05 for all experiments.

### Bioinformatic analyses

We utilized publicly available single-cell/nucleus RNA datasets to compare the similarity of human LGN to mouse LGN and to thalamocortical organoids^83,84,85^. To integrate these sequencing datasets, we utilized the scvi-tools package^86^. For data that were collected using SMART-seq, we normalized raw expression counts by gene length, as recommended in the scVI workflow. To normalize by gene length, we utilized gtftools^87^ to extract the gene lengths using the gtf files corresponding to the genome builds that the datasets were originally aligned to. Mouse genes were converted to human orthologs using Biomart, and those with a one-to-one match were retained. In the case the datasets contained experimental conditions, only wild-type conditions were retained. As some datasets differed in the cell types captured, we only compared neuronal types and relied on the original authors’ annotations for this. For larger datasets that exceeded 10,000 neurons, we further down-sampled randomly to a maximum of 10,000 neurons to ensure a more balanced dataset distribution.

To assess transcriptomic similarity amongst our datasets, we employed the reference similarity spectrum (RSS)^35,36^. Briefly, the RSS is a per-cell correlation-based metric that represents the similarity of query cells to an average reference expression profile. Using our integrated dataset, we split the data into reference and query, with the reference as the dissected lateral geniculate nucleus (LGN) single nucleus RNA-seq dataset from the Human Brain Cell Atlas^88^. The query was comprised of transcriptomic data collected from a secondary human dorsal LGN (dLGN) source^47^, mouse P56 dLGN2, mouse P3 dLGN3, mouse P98 OFB4, thalamic organoid6, and the thalamus cells from the human neural organoid cell atlas (HNOCA)^36^. RSS was calculated for each cell in the query datasets relative to the average expression profile of the 2000 most highly variable genes of the reference.

For cell-cell communication analysis, we performed cell-cell communication using LIANA+ with publicly available datasets^46^. For these analyses, we assessed interactions between organoid-derived RGCs^24^ with adult mouse SCN^89^, P3 mouse dLGN3, P56 mouse dLGN2, and P98 adult mouse olfactory bulb OFB^90^. Across all datasets, only the neurons were retained from wild-type conditions, using the original authors’ cell type annotations. For the SCN and OFB datasets, neurons were randomly down-sampled to 10,000 cells. We followed the LIANA+ steady-state ligand-receptor inference vignette, performing pairwise comparisons between the organoid RGC dataset and each target tissue, and utilized the consensus resource with the li.mt.rank_aggregate function. To ensure robustness, genes were required to be expressed in at least 10% of cells in a given cluster. Results were stratified by directionality to identify ’outgoing’ signaling (hRGC ligands targeting tissue receptors) and ’incoming’ signaling (tissue ligands targeting hRGC receptors). For the visualization of specific axon guidance interactions (e.g., CNTN2–CNTN1, L1CAM–FGFR2), interactions were compared across tissues using the aggregate rank score (lrscore).

## Acknowledgments

We thank Dr. Matthew N. Rasband for kindly sharing antibodies and advice, and Dr. Peter Bor-Chan Lin for assisting in the establishment of LGN and SCN mouse dissection techniques. We thank the Optical Imaging and Vital Microscopy Core at Baylor College of Medicine for technical support. This work was supported by the National Institutes of Health (NIH, grants R01EY033772, R01EY035254, R01EY032566, and R01EY030458 to M.A.S. and R01EY033022 and U24EY033269 to J.S.M., R01 EY036111 to N.M.T., and 5T32EY007001-47 in support of Claire Young), the Foundation Fighting Blindness (to M.A.S.), the TIRR foundation (to N.M.T), and the Cullen Foundation and Baylor Research Advocates for Student Scientists (to J.D.). This work was also supported by the RReSTORE Consortium (to K.H).

---

| Antibody | Type | Source | Catalog Number | ICC Dilution | Identifier |
| --- | --- | --- | --- | --- | --- |
| Ankyrin-G | Mouse Monoclonal | NeuroMab | 75-146 | 1:200 | RRID:AB_2877524 |
| βIV-spectrin | Rabbit polyclonal | Rasband lab | | 1:200 | RRID:AB_2315634 |
| BRN3 | Goat polyclonal | Santa Cruz | SC-6026 | 1:200 | RRID:AB_673441 |
| CHX10 (E-12) | mouse monoclonal | Santa Cruz | sc-365519 | 1:200 | RRID:AB_10842442 |
| FOXP2 | Sheep polyclonal | R&D system | AF5647 | 1:200 | RRID:AB_2107133 |
| GFAP | mouse monoclonal | Millipore-Sigma | G3893 | 1:200 | RRID:AB_477010 |
| GFP | Chicken polyclonal | Abcam | ab13970 | 1:2000 | RRID:AB_300798 |
| IBA1 | Goat polyclonal | Abcam | ab5076 | 1:100 | RRID:AB_2224402 |
| Ki67 | Rabbit Monoclonal | Invitrogen | MA5-14520 | 1:250 | RRID:AB_10979488 |
| MAP2 | Chicken polyclonal | Abcam | ab5392 | 1:2000 | RRID:AB_10842442 |
| MAP2 | Rabbit polyclonal | Proteintech | 17490-1-AP | 1:100 | RRID:AB_2137880 |
| NeuN | Rabbit polyclonal | Proteintech | 26975-1-AP | 1:100 | RRID:AB_2880708 |
| Pan sodium channel | Mouse Monoclonal | NeuroMab | 75-407 | 1:200 | RRID:AB_2877588 |
| SMI-312 | mouse monoclonal | BioLegend | 837904 | 1:500 | RRID:AB_2566782 |
| Synapsin-1 | Rabbit polyclonal | Millipore-Sigma | AB1543 | 1:200 | RRID:AB_2200400 |
| Synapsin-1 | Rabbit monoclonal | Cell Signaling | 5297 | 1:200 | RRID:AB_2616578 |
| tdTomato | Goat polyclonal | Origene | AB8181-200 | 1:500 | RRID:AB_3206272 |
| TUBB3 | Rabbit<br>Polyclonal | BioLegend | 802001 | 1:1000 | RRID:AB_2564<br>645 |
| VIP | Rabbit<br>polyclonal | Immunostar | 20077 | 1:1000 | RRID:AB_5722<br>70 |

## Supporting Information

**Supplemental Figure 1.**
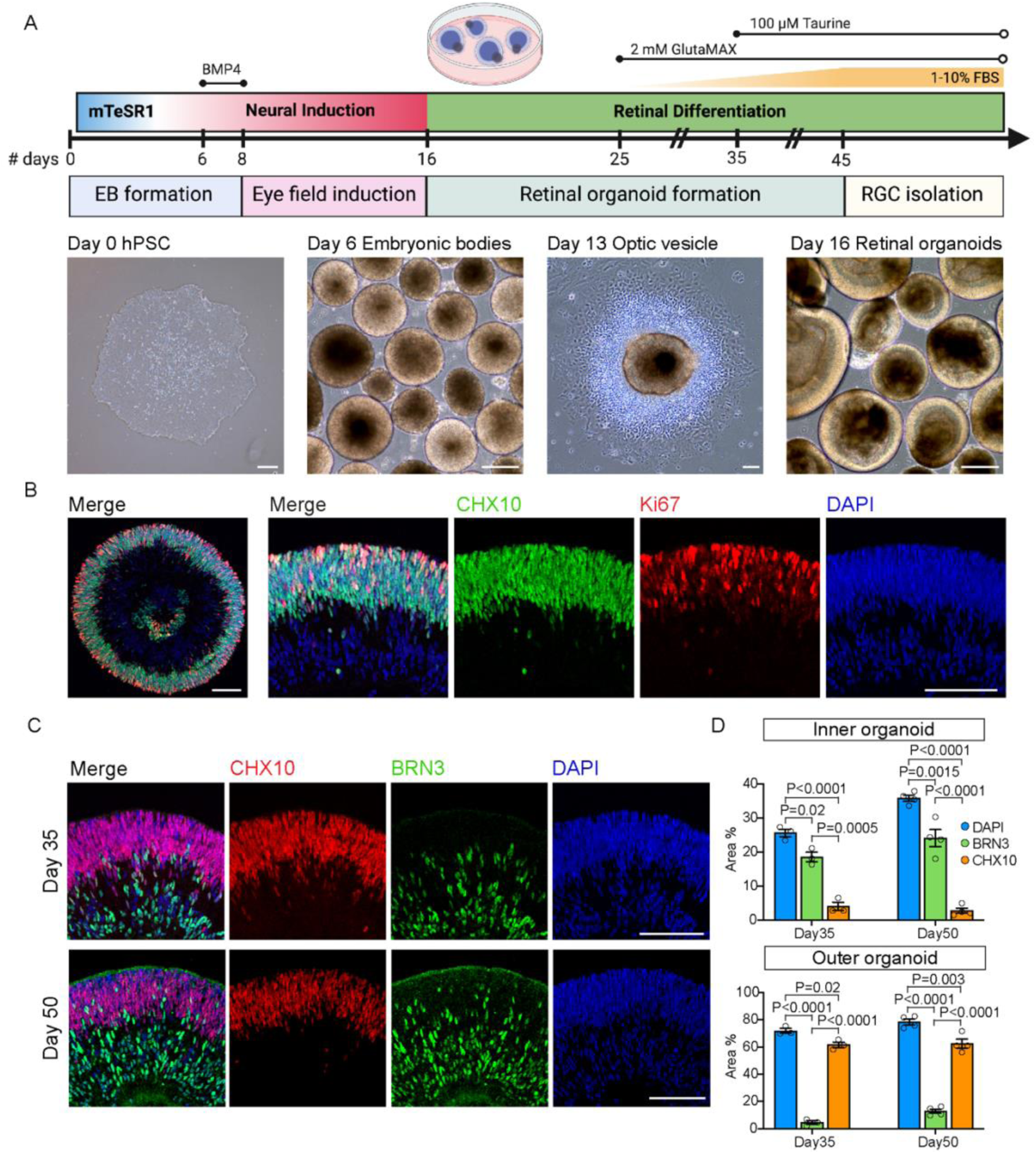
Generation of hRGCs from hPSC-derived retinal organoids. **A**. Schematic (upper panel) and representative images (lower panel) of the stages and the timeline for the induction of retinal organoids and RGC isolation. hRGCs were isolated from day 45 retina organoids for this study. Scale bars = 200µm. **B**. Cross section images of a day 35 retinal organoid shows that the organoids have a spherical structure with properly layered developing retinal progenitor cells (CHX10, green) in the outer retina that are actively cycling (Ki67, red). Fate committed cells migrate to the inner retina and are negative for these markers but positive for then nuclear marker DAPI (blue). Scale bars = 100µm. **C-D**. Representative cross section images (**C**) and quantification (**D**) comparing day 30 and day 50 retinal organoids. Developing human retinal organoids conserve spatial and temporal developmental patterns observed in the developing human eye. At day 30, subsets of fate committed RGCs (BRN3, green) can be observed developing within the inner retina, while retina progenitor cells remain in the outer retina (CHX10, red, **D**). By day 50, increased numbers of RGCs are observed in the inner retina, and the numbers of retinal progenitor cells are decreased (CHX10, red). n = 3 to 4 biological replicates for both day 35 and 50 retinal organoids. Scale bar = 100 µm. Data are presented as average values ± the s.e.m.

**Supplemental Figure 2:**
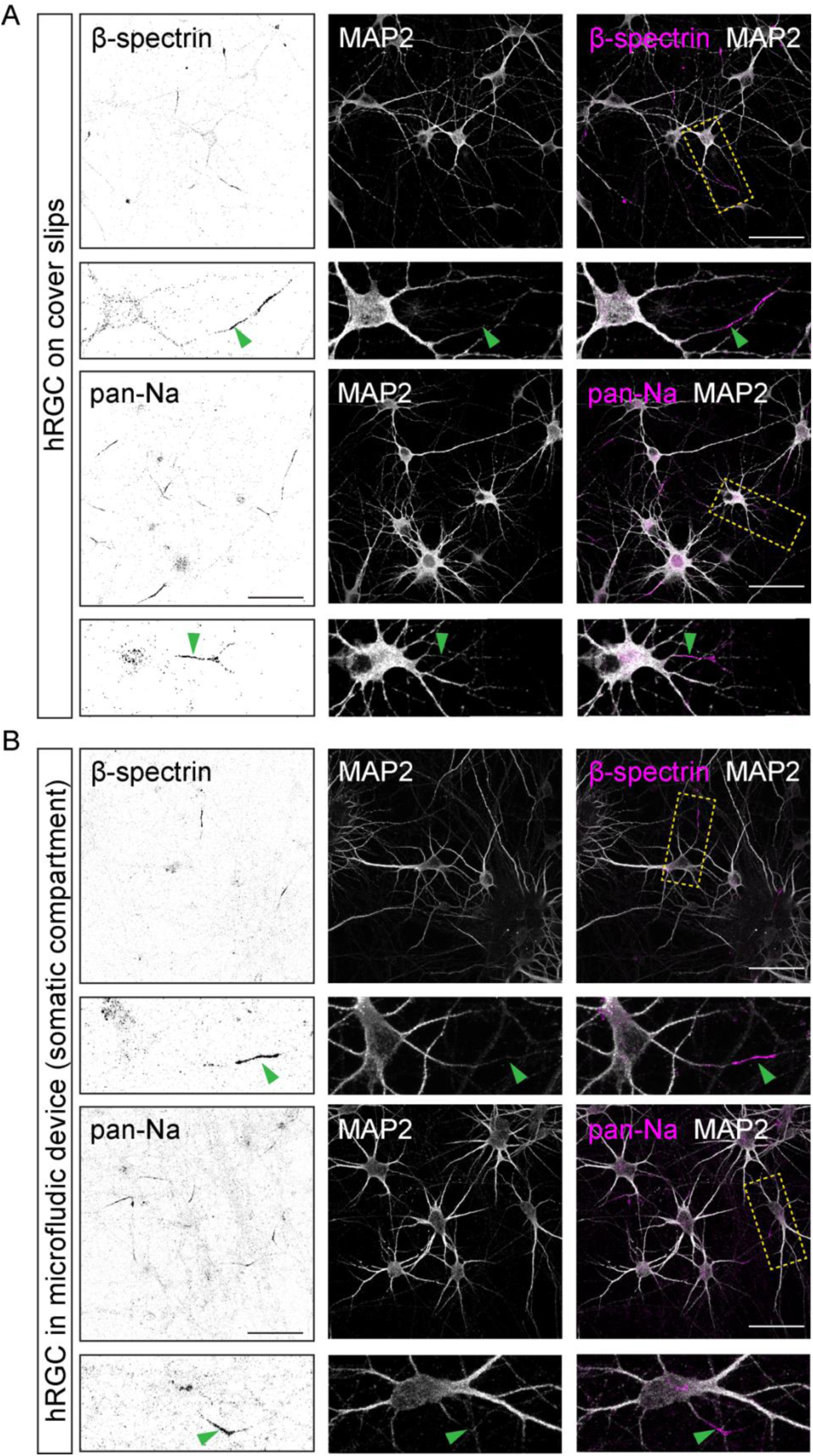
Differentiated hRGCs show additional axon initial segment localized proteins. **A-B.** Representative images from hRGCs in coverslips alone (**A**) or after culturing in microfluidics devices (**B**) for a period of 10 days following. Samples were stained for sodium channels (pan-Na, magenta upper panels) and β-Spectrin, both of which are known to be enriched in mature axon initial segments (AIS). In all cases, sodium channels and β-Spectrin were concentrated at AISs and were located proximal to the cell bodies of hRGCs (MAP2, white) both in coverslip conditions and in the microfluidics system (green arrows). Scale bars = 50 µm.

**Supplemental Figure 3:**
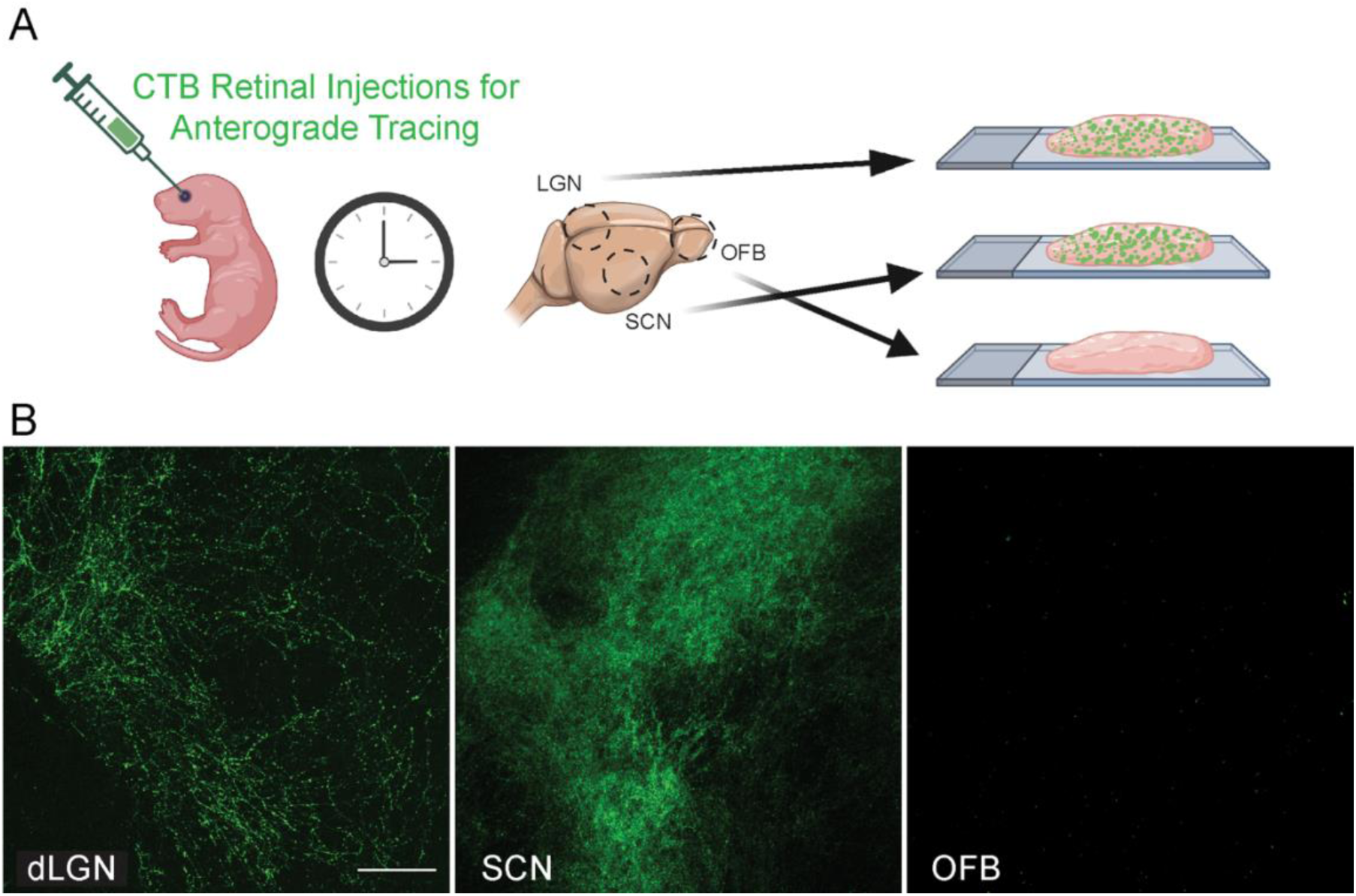
(**A**). Schematic of how CTB injections were administered and samples collected for imaging. P3 pups were intravitreously injected with CTB and brain regions from the LGN (**B**), SCN (**C**), and OFB (**D**) were collected 24 hours post injections. Scale bar: 100 um.

**Supplemental Figure 4.**
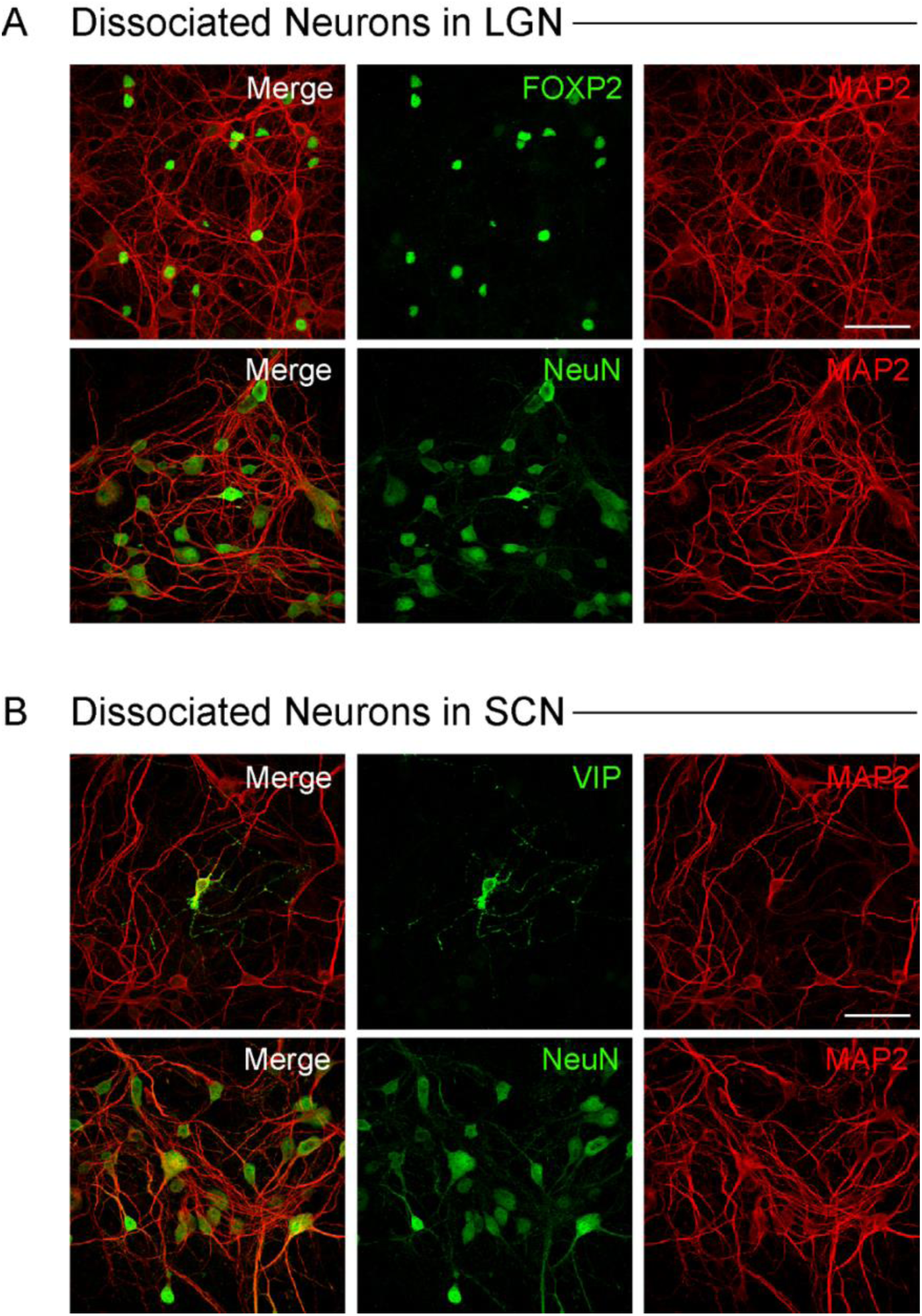
Cultured retinorecipient neurons express region-specific markers. **A**. Representative images of neurons in lateral geniculate nucleus (LGN) cultures following staining for FOXP2, a transcription factor enriched in a subset of LGN neurons (green, upper panel) or the neuron marker NeuN (green, lower panel) together with staining for the dendritic marker MAP2. A subset of LGN neurons were positive for FOXP2, suggesting that appropriate neural identity is maintained in these cultures. **B**. Representative images of neurons in suprachiasmatic nucleus (SCN) cultures following staining for vasoactive intestinal polypeptide (VIP), neurons essential for circadian entrainment (green, upper panel) or the neuron marker NeuN (green, lower panel) together with staining for the dendritic marker MAP2. Some SCN neurons were positive for VIP, suggesting that appropriate neural identity is also maintained in SCN cultures. Scale bars = 50 µm.

**Supplemental Figure 5.**
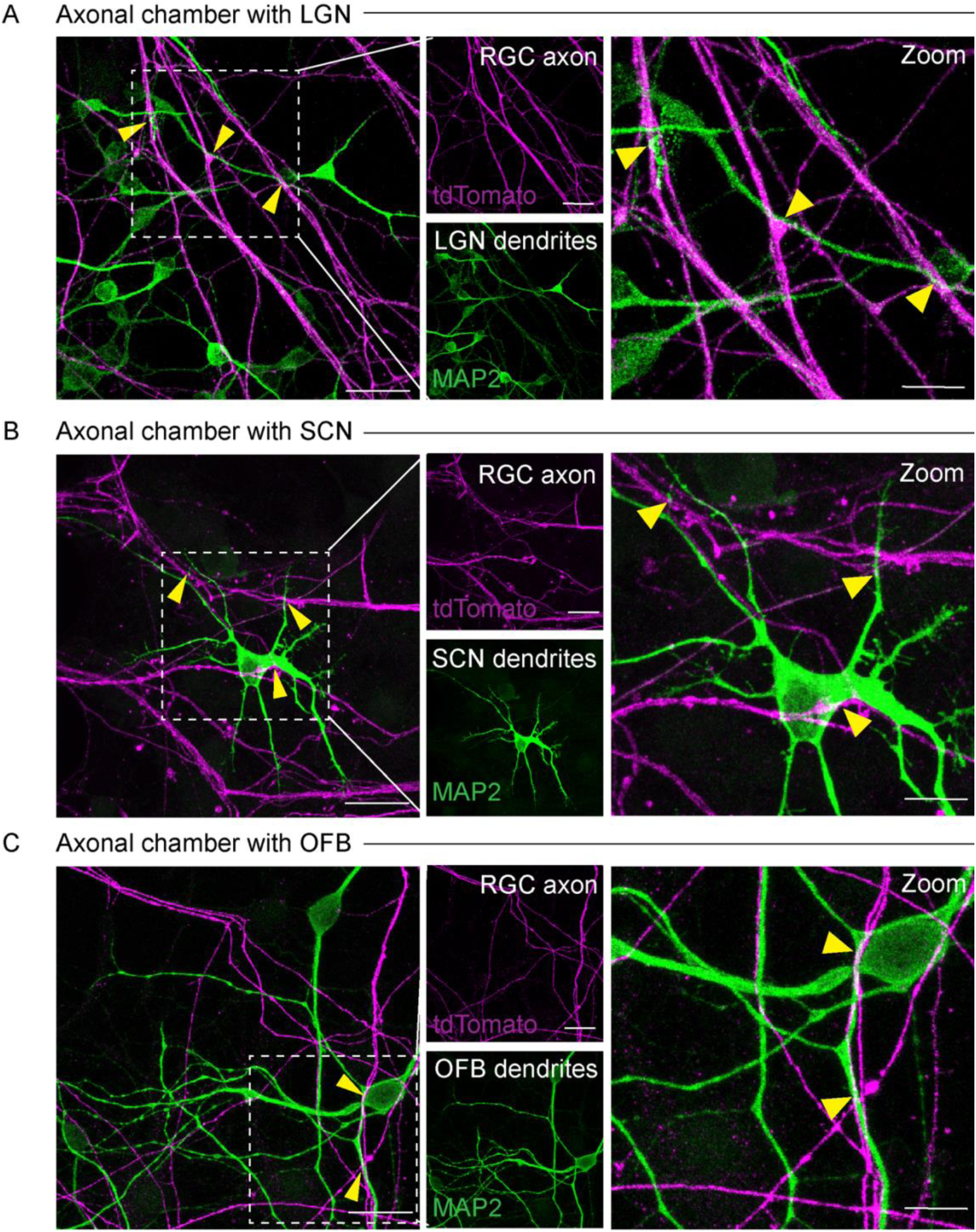
hRGC axons colocalize with postsynaptic target neurites in the axonal chambers. **A-C**. Representative low and high magnification (crop) images of the dendrite marker MAP2 (green) and tdTomato positive hRGCs axons (magenta) following imaging of axonal chamber of the addition of lateral geniculate nucleus (LGN), suprachiasmatic nucleus (SCN), or olfactory bulb (OFB) post-synaptic targets. In each case, hRGC axons co-localized and interfaced with the dendrites of post-synaptic neurons, suggesting the potential for synapse formation (yellow arrows). Scale bars = 25 µm.

**Supplemental Figure 6.**
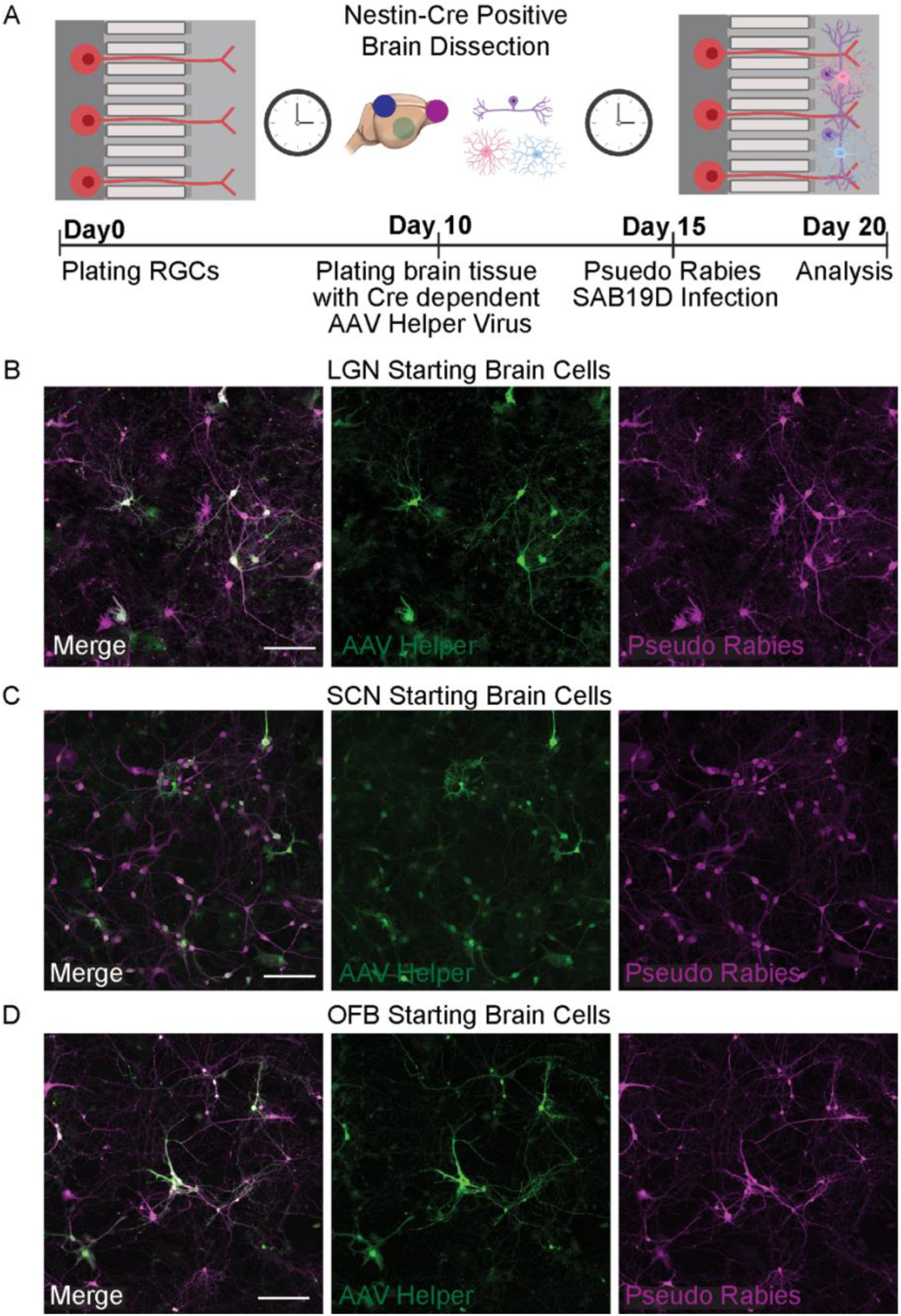
Cell transfection with AAV helper virus and Pseudo Rabies virus similar across different starter cells. **A**. Schematic of retrograde circuit tracing approach. hRGCs were grown in microfluidics chambers for 10 days. Following this period, dissociated cells from dissected control brain regions (OFB) and retinorecipient brain regions (LGN and SCN) were isolated from postnatal day (P) 0 to P3 mouse pups, infected with AAV-helper GFP,, and then plated in the axonal chamber. 5 days later, dissected brain regions were infected with the pseudo rabies virus. **B-D.** Representative images of the number of transfected cells from retinorecipient LGN and SCN neurons or OFB starter brain cells. Scale bars = 50 µm.

**Supplemental Figure 7:**
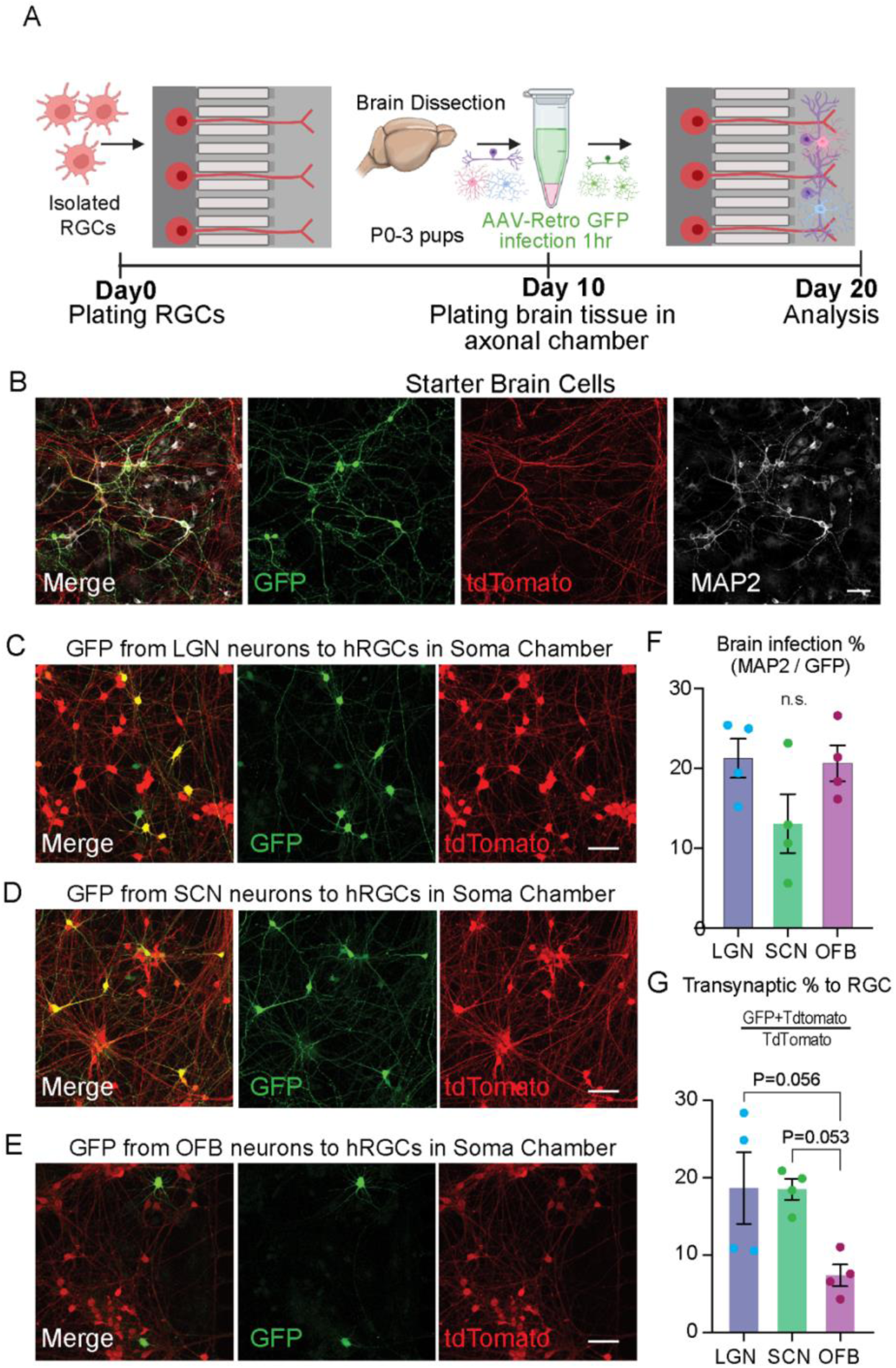
Retrograde labeling through AAV transfection. **A**. Schematic of retrograde circuit tracing approach. hRGCs were grown in microfluidics chambers for 10 days. Following this period, dissociated cells from dissected control brain regions (OFB) and retinorecipient brain regions (LGN and SCN) were isolated from postnatal day (P) 0 to P3 mouse pups, infected with AAV-Retro GFP, washed extensively, and then plated in then plated in the axonal chamber. GFP positive cells were quantified in the somatic chamber 10 days later. **B.** Representative images of AAV infection in the starter brain cell populations. Scale bars = 50 µm. **C-E.** Representative images of the number of transsynaptic labeled, GFP positive hRGCs (tdTomato+) following retrograde tracing from retinorecipient LGN and SCN neurons or OFB control neurons. Scale bars = 50 µm. **F-G** Quantification of AAV infection in the starter brain cells and transsynaptically labeled in hRGCs. No significant difference in AAV-Retro-GFP (green) was observed between the starter cell populations **(I)**. While a degree of connectivity was observed in all cases, LGN and SCN neurons tended to have higher degrees of connectivity relative to control OFB neurons (**J**). n = 4 tile images from 2 biological replicates, each image contains >200 neurons or hRGCs. Data are presented as averaged average values from 3 to 4 independent experiments ± the s.e.m.

**Supplemental Figure 8:**
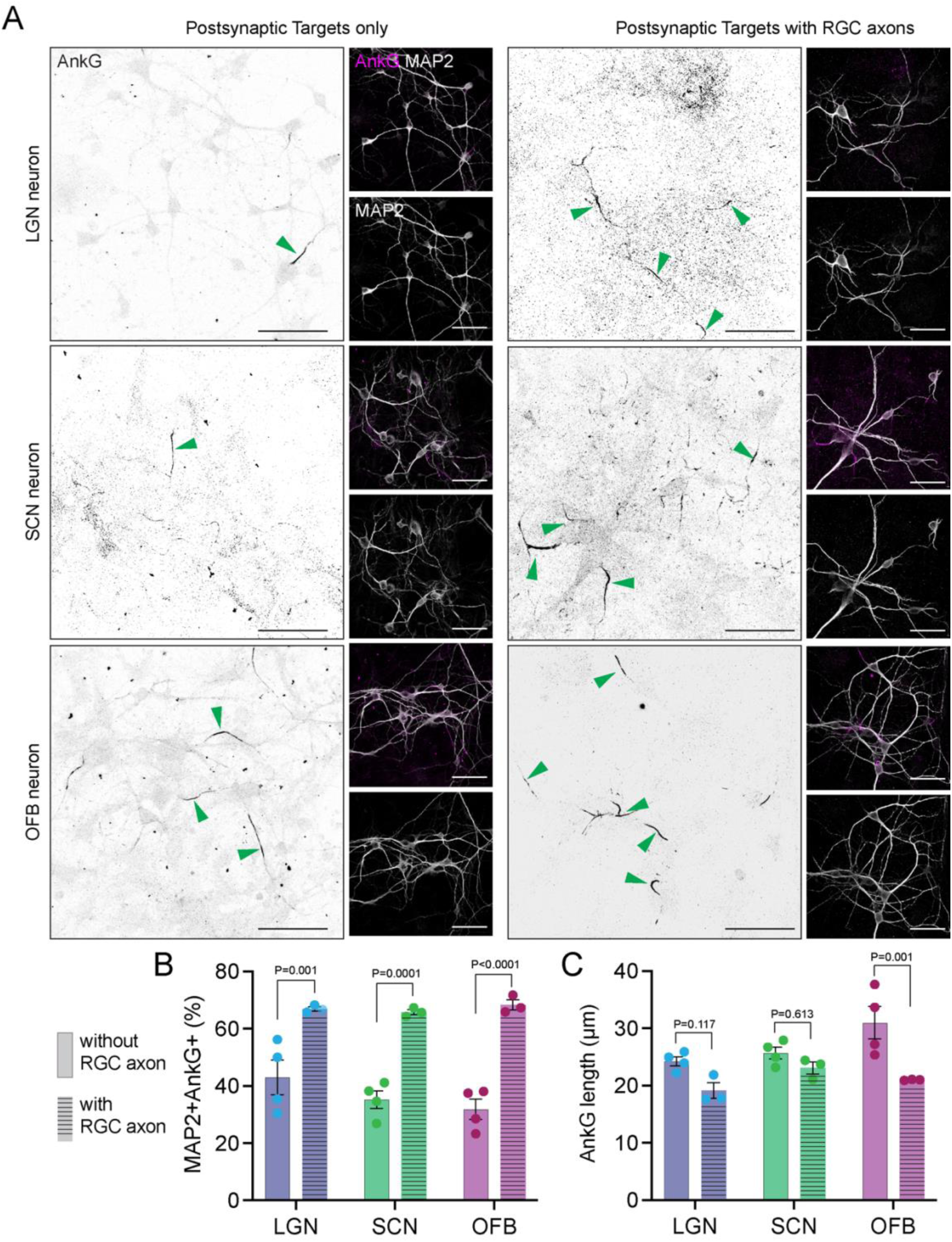
hRGCs promote the maturation of post-synaptic neurons in the microfluidics system. **A-C**. Representative low magnification and high magnification images of lateral geniculate nucleus (LGN), suprachiasmatic nucleus (SCN), or olfactory bulb (OFB) post-synaptic neurons plated in the microfluidics system grown in the absence and presence of hRGC axons following staining with the axonal initial segment structural protein AnkG (black/magenta) and the dendrite marker MAP2 (white). In each case, the presence of hRGC axons promoted the formation of additional AISs (green arrows). Scale bars = 50 µm. **B-C**. The number of postsynaptic neurons (MAP2+ cells) that contained an AIS (AnkG+) was significantly increased in all three target populations where neurons were cultured in the presence of pre-syanptic hRGC axons (**B**), while only OFB neurons showed a significantly longer AIS (**C**). n = 3 to 4 biological replicates. At least 4 images were used to quantify each replicate, and the number of hRGCs quantified was greater than 30 for each condition.

